# Multiomics characterization of the of the zoo-housed gorilla gut microbiome reveals bacterial community compositions shifts, fungal cellulose-degrading, and archaeal methanogenic activity

**DOI:** 10.1101/2022.11.15.516570

**Authors:** Isabel Mathilde Houtkamp, Martine van Zijll Langhout, Mark Bessem, Walter Pirovano, Remco Kort

## Abstract

We carried out a comparative analysis between the bacterial microbiota composition of zoo-housed western lowland gorillas and their wild counterparts through 16S rRNA gene amplicon sequencing. In addition, we characterized the carbohydrate-active and methanogenic potential of the zoo-housed gorilla microbiome through shotgun metagenomics and RNA sequencing. The zoo-housed gorilla microbiota showed increased alpha diversity in terms of bacterial species richness and a distinct composition from that of the wild gorilla microbiota, including a loss of abundant fiber-degrading and hydrogenic Chloroflexi. Metagenomic analysis of the CAZyome indicated predominant oligosaccharide-degrading activity, while RNA sequencing revealed diverse cellulase and hemi-cellulase activities in the zoo-housed gorilla gut, contributing to a total of 268 identified carbohydrate-active enzymes. Metatranscriptome analysis revealed a substantial contribution of 38% of the transcripts from anaerobic fungi and archaea to the gorilla microbiome. This activity originates from cellulose-degrading and hydrogenic fungal species belonging to the class Neocallimastigomycetes, as well as from methylotrophic and hydrogenotrophic methanogenic archaea belonging to the classes Thermoplasmata and Methanobacteria, respectively. Our study shows the added value of RNA sequencing in a multiomics approach and highlights the contribution of eukaryotic and archaeal activities to the gut microbiome of gorillas.

## Introduction

Gut microbes are necessary for mammals to digest complex polysaccharides in plants (fibers), as the microbial genomes encode the metabolic potential to support the hydrolysis and fermentation of plant material (Gomez, 2014). The gut microbiome has even allowed the evolution of herbivory as a dietary niche, and many mammals now rely completely on plant material as a dietary source (Moeller & Sanders, 2020). After millions of years of co-evolution, the gut microbiome is not only embedded in host metabolism, but also in the immune and endocrine systems, with hosts and microbiomes shaping each other’s genomes (Moeller et al., 2016). The host microbiome can rapidly adapt to composition and function in response to dietary and other lifestyle changes. Although all mammals may have experienced such changes, the most frequent and drastic lifestyle transitions have occurred in human history, with industrialization being the most recent and fast-paced example. Human exposure to microbes has been drastically altered in industrialized societies owing to antibiotics, higher sanitation levels, non-emergency (planned) C-sections, formula infant feeding, and processed foods, which are low in dietary fiber (Sonnenburg and Sonnenburg, 2019a; Sonnenburg and Sonnenburg, 2019b). Comparisons between industrialized microbiomes and those of hunter-gatherer societies indicate that, while the hunter-gatherer microbiome has a rich diversity of species and carbohydrate active enzymes, the industrialized microbiome has a lower taxonomic diversity and is enriched with mucin-degrading enzymes. An increased abundance of mucin-degrading genes is a sign of low nutrient availability for the gut microbiome, allowing host mucin-degrading microbes to flourish. While the microbiome may have adapted to industrialization, concerns have been raised about host biology because of the ‘asymmetric plasticity’ between the rapidly adapting microbiome and the relatively stable mammalian genome. Sonnenburg and Sonnenburg (2019b) postulated that because of rapid lifestyle transitions, the microbiome can become incompatible with its host, possibly leading to the emergence of noncommunicable chronic diseases in industrialized societies.

Interestingly, the effects of industrialization on the microbiome also apply to zoo-born primates. Indeed, zoo-housed primates are also exposed to higher sanitation levels, antibiotics, sometimes formula infant feeding, and most consistently, to a diet lower in plant complex carbohydrates than their wild counterparts. Efforts have been made to understand the effects of zoo-housing on the mammalian microbiome to gain a general understanding of lifestyle effects, improve the health of zoo-housed animals, and support conservation efforts. Clayton et al. (2016, 2018) found that the microbiota of zoo-housed primates (doucs, *Pygathrix nemaeus*, and mantled howler monkeys, *Alouatta palliata*) was reduced in diversity and enriched in genera that dominated the microbiota of industrialized humans, such as *Bacteroides* and *Prevotella*. According to observations in North American Zoos, the gut microbiomes of chimpanzees, bonobos, and gorillas contain 30% fewer genera than those of their wild fellow species, which is similar to the loss of genera in industrialized humans compared to humans living a hunter-gatherer lifestyle (Nishida & Ochman, 2021). In contrast, Campbell et al. (2020) showed that zoo-housed gorillas had almost two times higher amplicon sequencing variant (ASV) richness than wild gorillas, while Shannon diversity values were similar between the two groups. A recent study by Narat et al. (2020) confirmed a significantly higher microbial richness in zoo-housed gorillas than in wild gorillas, although Shannon diversity values also increased. Despite this, the microbiota of zoo-housed apes is still most similar to that of wild apes and more similar to that of non-industrialized humans than Westernized humans. This is attributed mainly to the presence of *Treponema* species, which are rarely present in industrialized human microbiomes but are common members of the hunter-gatherer microbiome and zoo-housed and wild great apes (Campbell et al., 2020).

However, species that are considered characteristic of the wild great ape microbiome are often lost in the zoo-housed microbiomes (Campbell et al. 2020; Clayton et al. 2016). The effects of lifestyle and diet on microbiome diversity and composition are evident in both humans and non-human primates. In gorillas, this is visible to the naked eye. The abdomens of wild gorillas are bloated, a result of fermentation of plant material by the microbiome leading to gas production by methanogens. The abdomens of zoo-housed gorillas often appear flat, indicating low levels of fermentation of plant material (Masi, 2011). Wild gorillas are herbivores that rely strongly on the gut microbiome to degrade plant material. The microbiota of western lowland gorillas is characterized by a high relative abundance of the phylum Chloroflexi compared to other great apes and humans and a high abundance of the phylum Spirochaetes, which is only present in high abundance among other primates in chimpanzees (Hicks et al., 2018).

Most data on the non-human primate microbiome originate from 16S rRNA gene amplicon sequencing and are thus compositional in nature. In human microbiome studies, shotgun metagenomics is now widely used for taxonomic and functional microbiome characterization (Zhang et al., 2019). For gorillas, such data are still limited; previous studies utilized shotgun metagenomics to study the prevalence of antibiotic-resistance genes in non-human primate microbiomes (Campbell et al., 2020), or to show seasonal fluctuations in the abundance of functional pathways in the great ape microbiome (Hicks et al., 2018). Whereas metagenomic studies can identify the potential of carbohydrate-active enzymes in the gut microbiome, metatranscriptomics can reveal which genes are expressed under certain conditions (Zhang et al., 2019), and thus reveal the functional activity of the microbiome. To the best of our knowledge, metatranscriptomics has never been used to study the microbiome of gorillas. Here, we studied the compositional differences between the gut microbiota of wild and zoo-housed gorillas by combining the 16s rRNA gene amplicon sequencing data of the gorilla population in ARTIS (the Amsterdam Royal Zoo) with previously published 16S rRNA gene amplicon sequencing data from wild gorillas (WG) and zoo-housed gorillas (ZHG) (Narat et al., 2020; Campbell et al., 2020). Furthermore, we present a metagenomic and metatranscriptomic analysis of the microbiome of the dominant male (‘silverback’) of the ARTIS gorilla population. We report the composition, fiber-degrading potential, and activity of the ZHG microbiome.

## Methods

### Diet of ARTIS zoo-housed gorillas

The ARTIS gorillas were daily fed gorilla pellets (Marion Leaf Eater Food, Plymouth, USA), 0.5 – 1% of their body weight. They received a wide array of vegetables that differed each day (rotated weekdays), including celeriac, endive, fennel, tomatoes, chicory, carrots, and a handful of nuts, seeds, rice, and muesli per animal. Tree branches of the willow tree and, when available, herbs from the ARTIS garden were added daily. An egg or tofu was offered once or twice a week. Hay was always available in the enclosure.

### Comparative analysis of zoo-housed and wild gorilla microbiota

We utilized previously published 16S rRNA gene V4 and V3-V4 amplicon sequencing data (Illumina technology) of WG and ZHG fecal samples obtained by Campbell *et al*. (2020) and Narat *et al*. (2020) for a comparative analysis in terms of diversity and composition with the microbiota data of ZHG fecal samples obtained in this study. In the study by Campbell et al. (2020), fecal samples from 28 WG’s were collected in Nouabalé-Ndoki National Park in the Republic of the Congo in February and March 2013. This period corresponded to the transition period between the dry and wet seasons and was a period of lowered dietary diversity. In addition, fecal samples from 15 ZHG’s were collected in 2017 in two zoos in the United States of America: four in the St. Louis Zoo (St. Louis, MO) and 11 in the Lincoln Park Zoo (Chicago, IL). Narat *et al*. (2020) collected 15 fecal WG samples from a forest site in southeastern Cameroon (between the towns of Yokadouma and Moloundou) in January 2016. This period corresponds to the dry season, which is characterized by low fruit availability and consumption of high-fiber fallback food. Additionally, fecal samples from the six ZHG’s were collected from a European zoo site in November 2017. A more detailed description of these datasets can be found in Campbell *et al*. (2020) and Narat *et al*. (2020). The 16S rRNA gene amplicon sequence data, generated with an Illumina MiSeq instrument using paired-end 2 × 250 bp and 2 × 270 bp protocols, respectively, were retrieved from the Sequence Read Archive (accession numbers: PRJNA539933 and PRJNA666756).

### Collection of fecal samples

A total of 15 fecal samples for 16S rRNA gene V3-V4 amplicon sequencing were collected from the ARTIS gorilla population between August 2019 and the end of January 2020. In July and August 2021, additional fecal samples from the silverback were collected in duplicates at two time points. Samples were collected within 30 min of defecation and frozen at −60°C. These four samples were subjected to shotgun metagenomics and RNA-seq analyses.

### DNA and RNA extraction, library preparation and Illumina sequencing

DNA was extracted from gorilla feces using the ZymoBIOMICS™ 96 MagBead DNA Kit (Zymo Research). DNA QC quantitation was performed with Quant-iT™ dsDNA Broad-Range Assay Kit (Invitrogen) and agarose gel electrophoresis for DNA integrity. The DNA served as a template for PCR amplification of a part of the 16S rRNA gene and subsequent sequencing using an Illumina MiSeq instrument. In short, amplicons from the V3-V4 regions of the 16S rRNA gene were generated by PCR using primers 341F and 785R, which generated a 444 bp amplicon (Klindworth et al., 2013) complemented with standard Illumina adapters. Index Primers were added to the amplicons of each sample through a second PCR cycle. The PCR products were purified using Agencourt© AMPure^®^ XP (Beckman Coulter), and the DNA concentration was measured by fluorometric analysis (Quant-iT™, Invitrogen). Subsequently, PCR amplicons were pooled using equimolar quantities, followed by sequencing on an Illumina MiSeq instrument using a paired-end 2 × 300 bp protocol and indexing. For samples subjected to shotgun metagenomics and RNA-seq, the ZymoBIOMICS™ ZymoD4300 kit was used to extract DNA and the ZymoBIOMICS™ kit Zymo R2040 to extract RNA. Library preparation was performed using Illumina RNA and DNA library preparation kits (20037135, 20020595, 20022371, and FC-131-1096). RNA integrity was assessed using a Bioanalyzer system (Agilent RNA 6000 Nano kit art5067-1511). RNA quality was ensured through the RIN value in combination with manual evaluation of the traces. Sequencing was performed on an Illumina NovaSeq 6000 instrument (paired-end, 2 × 150 bp) using an SP and S2 flow cell. Reads containing phiX control signals were removed using an in-house filtering protocol (Baseclear BV, Leiden, the Netherlands). In addition, reads containing (partial) adapters were clipped to a minimum read length of 50 bp. The second quality assessment was based on the remaining reads, using the FASTQC quality control tool version 0.11.8 (Andrews, 2010).

### Analysis of 16S rRNA gene amplicon sequencing data

MiSeq paired-end sequence reads from Campbell et al. (2020), Narat et al. (2020), and our own study were subsampled to 10 MBp, corresponding to 20.000, 18.518/18.519, and 16.628-16.821 reads, respectively (rarefaction curve, Supplemental Figure S1). For Narat et al. (2020) and our own study, the 16S rRNA gene V3-V4 amplicons were trimmed to retain only the V4 region, allowing for a consistent comparison with the samples of the Campbell *et al*. (2020) study. Hereafter, the paired-end sequences were collapsed into pseudoreads (reads that have been merged based on their overlapping sequences) using sequence overlap with USEARCH version 9.2 (Edgar & Bateman, 2010). Classification of these pseudoreads was based on the results of alignment with SNAP version 1.0.23 against the RDP database version 11.5 for bacterial organisms (Cole et al., 2014). OTU clusters were generated at 97% similarity with the USEARCH function “cluster_otus” based on pseudoreads. Reads that could not be assigned to sequences originating from the Domain of Bacteria were excluded. To analyze beta diversity, sequence counts were transformed to relative abundances, and Bray-Curtis distances were calculated for Principal Coordinate Analysis (PCoA). PCoA significance was tested using a non-parametric multivariate analysis of variance (ADONIS). Alpha diversity metrics were calculated from untransformed genus-level counts, and the significance of differences in alpha diversity metrics between groups was determined using the Wilcoxon rank-sum test LEfSE (Segata et al., 2011) was performed for differential abundance analysis between wild and zoo-housed samples using default CPM normalization on taxa counts. Genera with an LDA score of at least 4 and a p-value below 0.05 were considered significantly enriched. The above-mentioned analyses and visualizations were carried out with R studio (R version 4.1.3) using the following packages: phyloseq 1.38.0 (McMurdie & Holmes, 2013), vegan 2.6-2 (Dixon, 2003), metacoder 0.3.5 (Foster et al., 2017) and microbiomeMarker 1.2.1 (Cao et al., 2022).

### Taxonomic assignment of DNA and RNA reads

Kraken2 (2.1.1)/Bracken (2.6.1) (Wood et al., 2019; Lu et al., 2017) was used to estimate the taxonomic composition of the metagenomes and metatranscriptomes using the Kraken/Braken database, which is based on the RefSeq of the NCBI (feb-2022). We extended this database with the addition of 6 Neocallimastigomycetes genomes: *Anaeromyces sp. S4 v1*.*0* (GenBank: GCA_002104895.1), and *Neocallimastix sp. G1 v1*.*0* (GenBank: GCA_002104975.1), *ASM1694683v1* (GenBank: GCA_016946835.1), *Orpinomyces* sp. strain C1A (GenBank: GCA_000412615.1), *Piromyces* sp. finnis v3.0 (GenBank: GenBank: GCA_002104945.1) and *Piromyces sp*. E2 v1.0 (GenBank: GCA_002157105.1). Assignments to the order of primates were filtered out, as these were considered to be contamination, and total sum scaling was applied.

### Functional profiling of metagenome and metatranscriptome

Shotgun metagenomic reads were filtered for polyG tails, a known artifact of Illumina NextSeq two-color dye technology (fastp version 0.23.1 (Chen et al., 2018) using the default -g option (polyG tail length = 10)). In addition, we used fastp to remove low-quality reads using the default -q option (bases with a phred quality > 15 were considered qualified) and -u option (filter reads with more than 40% unqualified bases). Metagenome and metatranscriptome reads were filtered for host cDNA using STAR aligner version 2.7.9a (Dobin et al., 2013) and Kamilah_GGO_v0 reference genome (Refseq GCF_008122165.1). The RNA-seq reads were subjected to the same preprocessing, but additionally filtered for ribosomal RNA with sortmerna version 4.3.4 (Kopylova et al., 2012) by aligning reads against all SILVA databases of release 138.1, which contain small (16S/18S, SSU) and large (23S/28S, LSU) ribosomal subunits of Archaea, Bacteria and Eukarya (Quast et al., 2013). Preprocessed DNA and RNA reads were annotated using HUMAnN3.0, version 3.0.0, (Beghini et al., 2021), with UniRef90 (version 20191b_full) as a reference database. To increase mapping rates, reads were assembled with MEGAHIT (1.2.9) (Li et al., 2015), genes were clustered using CD-HIT (4.8.1) (Fu et al., 2012), annotated with Prokka (v1.14.5) (Seemann, 2014), and EggNOG v. 5.0 (Huerta-Cepas et al., 2019) using the EggNOG-mapper (v2.0.0) (Cantalapiedra et al., 2021). Subsequently, the original preprocessed reads were realigned against the deduplicated contigs with bowtie2 v2.4.2 (Langmead & Salzberg, 2012) and subjected to a second analysis with HUMAnN3.0.0, using the contig KEGG and COG annotation as custom mapping files, to obtain relative abundances of these functional groups. Normalization was performed using HUMAnN3’s normalization function. Analyses were executed with Nextflow version 21.10.5 (build 5658) (di Tommaso et al., 2017) for reproducibility.

### Annotation of Carbohydrate Active Enzymes

To quantify the relative abundances of carbohydrate-active enzyme (CAZyme) families, we ran HUMAnN3.0, on the read-contig alignments with a custom id-mapping file containing the corresponding CAZyme family for each contig. To create this file, clustered contigs were aligned to the dbCAN2 version of the CAZy database (version 10: CAZyDB.09242021). fa) using run_dbcan (v2.0.11) (Zhang et al., 2018), employing the option for alignment with HMMER in metagenome mode. Normalization was performed using the recommended normalization function of HUMAnN3.0.

### Detection of fungal genes and transcripts

After preprocessing, ShortBRED (v0.9.4) (Kaminski et al., 2015) was used for sensitive detection of proteins of interest. We created markers for several proteins originating from Neocallimastigomycetes species downloaded from UniProt (release 2022_01). Preprocessed DNA and RNA reads were searched against several markers for proteins of interest. A first set of markers contained all Neocallimastigomycetes sequences in UniProt annotated with the subcellular localization ‘hydrogenosomal’. A second set of markers consisted of all malic enzyme sequences (malate dehydrogenase [decarboxylating] EC 1.1.1.39) assigned to Neocallimastigomycetes and the latest sets of GH48 and GH6 proteins. Clustid was set to 85% and the minimum marker length to 8 amino acids, based on the authors of ShortBRED’s own parameter optimization, except for the marker set of malic enzymes, for which the clustid option was set to 100% because of high sequence similarity. A workflow overview of the shotgun metagenomics and RNA-seq data analysis is shown in Supplemental Figure S2.

## List of abbreviations

CAZyome: collective of carbohydrate-active enzymes
MG: metagenome
MT: metatranscriptome
RNA-seq: RNA sequencing
WG: wild gorilla
ZHG: zoo-housed gorilla
ACE: abundance-based coverage estimators
GH: glycoside hydrolase

## Ethics approval

This study was approved by the ARTIS Amsterdam Royal Zoo Ethics Committee. All methods were noninvasive and part of routine preventive health monitoring.

## Data availability

All sequence data on fecal samples of the ARTIS gorilla population have been deposited in the Sequence Read Archive (SRA) under BioProject number PRJNA827108. All scripts used for data analysis are available in a dedicated GitHub repository (Houtkamp, 2022).

## Results

### Beta-diversity of bacterial communities differs between zoo-housed and wild gorillas

In this study, we compared the composition of zoo-housed gorilla microbiota to that of wild gorillas. To accomplish this, we combined previously published amplicon sequencing data (Campbell et al., 2020; Narat et al., 2020) from wild and zoo-housed western lowland gorillas with our amplicon sequencing data from 15 fecal samples from a group of Western lowland gorillas in ARTIS, the Amsterdam Royal Zoo. Principal Coordinate Analysis (PCoA) based on the Bray-Curtis distance between samples indicated that bacterial community compositions from different studies clustered according to whether they originated from zoo-housed or wild gorillas (Figure 1). Zoo-housed gorilla samples from the three different studies converged and were clearly separated from wild gorilla samples. Non-parametric multivariate analysis of variance (ADONIS) calculated for the Bray-Curtis distances between zoo-housed gorilla and wild gorilla samples revealed that intra-group variability was lower than inter-group variability (*p = 0*.*001*), indicating that the microbiota composition of zoo-housed gorillas was significantly different from that of wild gorillas.

**Figure 1.**
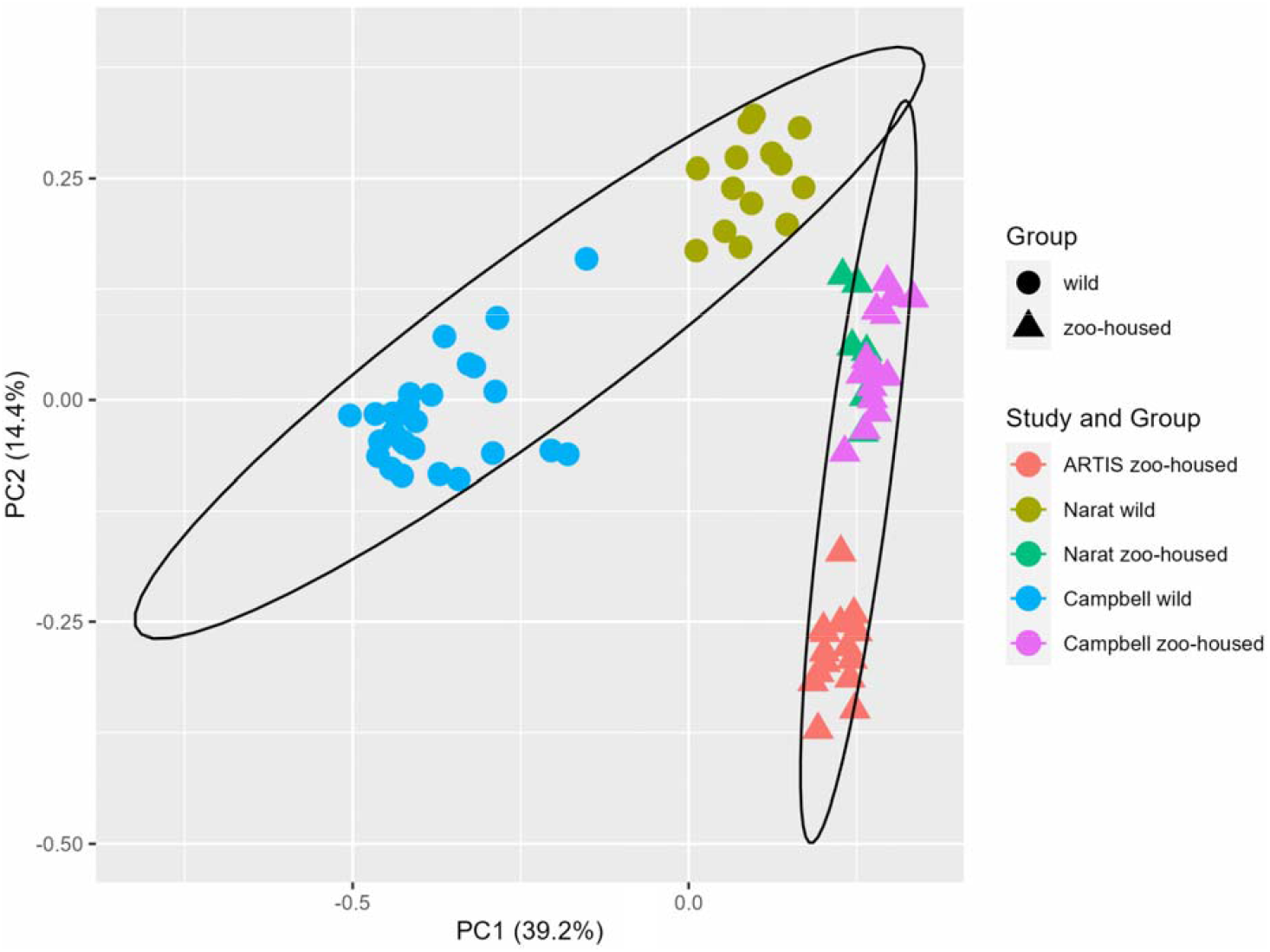
Principal Coordinate Analysis based on Bray-Curtis dissimilarities. The first two principal coordinates showed that, combined, they explain 53.6% of the observed variation in the zoo-housed and wild gorilla gut microbiota based on 16S rRNA gene V4 amplicon sequences. Colors indicate the study groups from which the samples originated. To aid interpretation, shapes indicate whether samples originated from wild or zoo-housed gorillas, regardless of the study.

### Microbiota alpha-diversity is elevated in zoo-housed gorillas

Alpha diversity indices revealed a significant increase in genus-level bacterial diversity in zoo-housed gorilla samples compared to wild gorilla samples with respect to estimated richness (Chao1 index, *p* = *0*.*0001*), abundance-based coverage estimator (ACE) index (*p* < 0.0001), and Fisher diversity index (*p* < 0.0001). No significant difference was found in the Shannon index between the zoo-housed gorilla and wild gorilla samples (*p* = 0.13) (Figure 2). Chao1, ACE, and Fisher’s exact test are known to be sensitive to species richness, whereas Shannon’s diversity index also reflects species evenness. Therefore, we concluded that microbiota richness was higher in the zoo-housed gorilla samples than in the wild gorilla samples.

**Figure 2.**
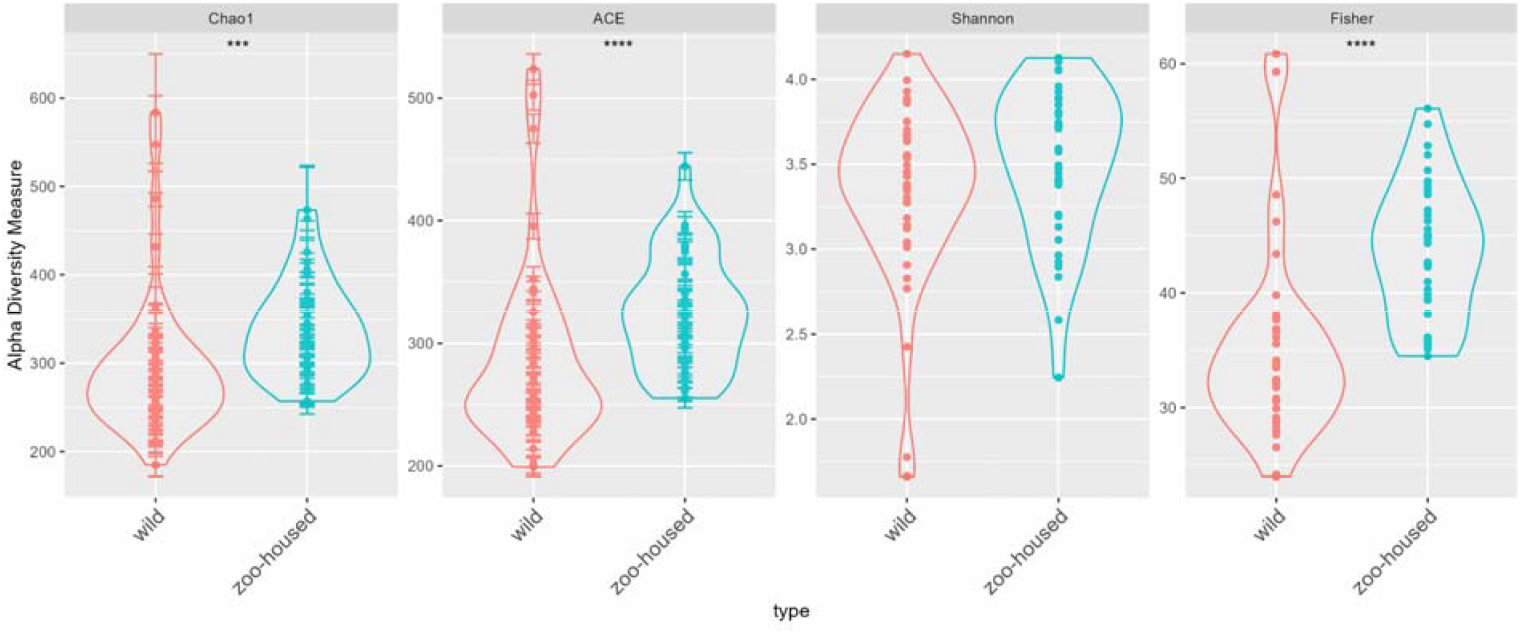
Microbiota alpha diversity of wild and zoo-housed gorillas. Alpha diversity metrics were calculated at the genus level for the zoo-housed and wild gorilla gut microbiota, based on 16S rRNA gene V4 amplicon sequences. Alpha diversity metrics for wild gorillas (red) and zoo-housed gorillas (blue-green). The diversity indices included **A)** Chao1, **B)** abundance-based coverage estimator index (ACE), **C)** Shannon, and **D)** Fisher. The indicators of significance levels **** = p < 0.0001 and *** = p < 0.001 were computed using Wilcoxon rank sum tests.

### Characteristic members of the wild gorilla bacterial microbiota vanish in zoo-housed gorillas

At the phylum level, the wild gorilla samples were, in order of abundance, dominated by Firmicutes, Chloroflexi, Bacteroidetes, Actinobacteria, Proteobacteria, and Spirochaetes (Supplemental Table S2). The most striking difference was the reduction in Chloroflexi, from 15.1% in wild gorillas to 0.13% in zoo-housed gorillas (Supplemental Figure S4). In zoo-housed gorillas from ARTIS (this study) and those described by Narat et al. (2020), Chloroflexi could not be detected at all, whereas they were detectable at low levels in zoo-housed gorillas, as described in the study by Campbell et al. (2020) (Supplemental Figure S4). At the genus level, the dominant taxa differed greatly between the zoo-housed gorillas and wild gorillas. An overview of the top ten genera in both the zoo-housed gorillas and wild gorilla samples and their relative abundances is provided in Supplemental Table S1; only the genera *Prevotella* and *Clostridium* were present in the top ten of both groups. The per-sample Z-transformed relative abundances are shown in the heatmap in Figure 3.

**Figure 3.**
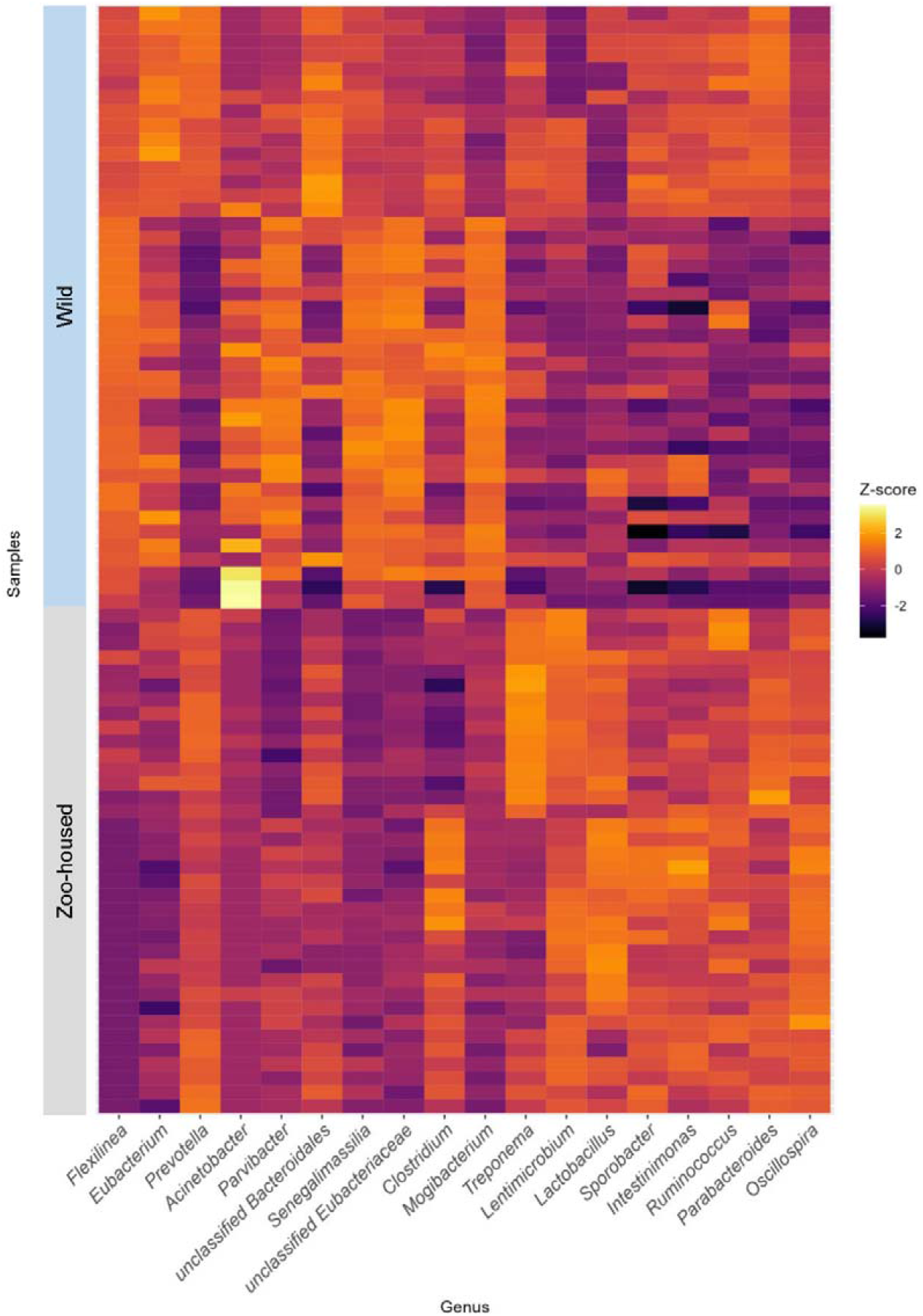
Heatmap depicting the top 10 dominant bacterial genera in WG and ZHG. Relative abundances have been Z-transformed and from left to right the top 10 genera of WG and the top 10 genera of ZHG samples have been selected for display. As two of the bacterial genera were in the top 10 of both groups, this combined heatmap shows the abundance per sample for 18 genera.

Linear discriminant analysis effect size (LEfSe) was used to identify specific enriched taxa in the gut microbiota of wild and zoo-housed gorillas (Figure 4). Wild gorilla samples were significantly enriched for nine bacterial genera, including *Flexilinea* (Sun et al., 2016b), in line with the above-mentioned lack of detection of species from the phylum Chloroflexi in zoo-housed gorilla samples. This reduction in Chloroflexi in zoo-housed gorilla samples is a striking observation, since Chloroflexi are considered a characteristic species of the native gorilla microbiota that degrade carbohydrates in high-fiber fallback foods consumed during dry seasons (Hicks et al., 2018). A second characteristic taxon of the native gorilla microbiota that was reduced in zoo-housed gorilla samples was *Olsenella* (phylum Actinobacteria). However, the characteristic genus *Treponema* was preserved in zoo-housed gorilla samples and was found to be enriched. The oligosaccharide-consuming genera *Lactobacillus* and *Lentimicrobium* (Sun et al.,2016a) were also enriched in the zoo-housed gorilla samples.

**Figure 4.**
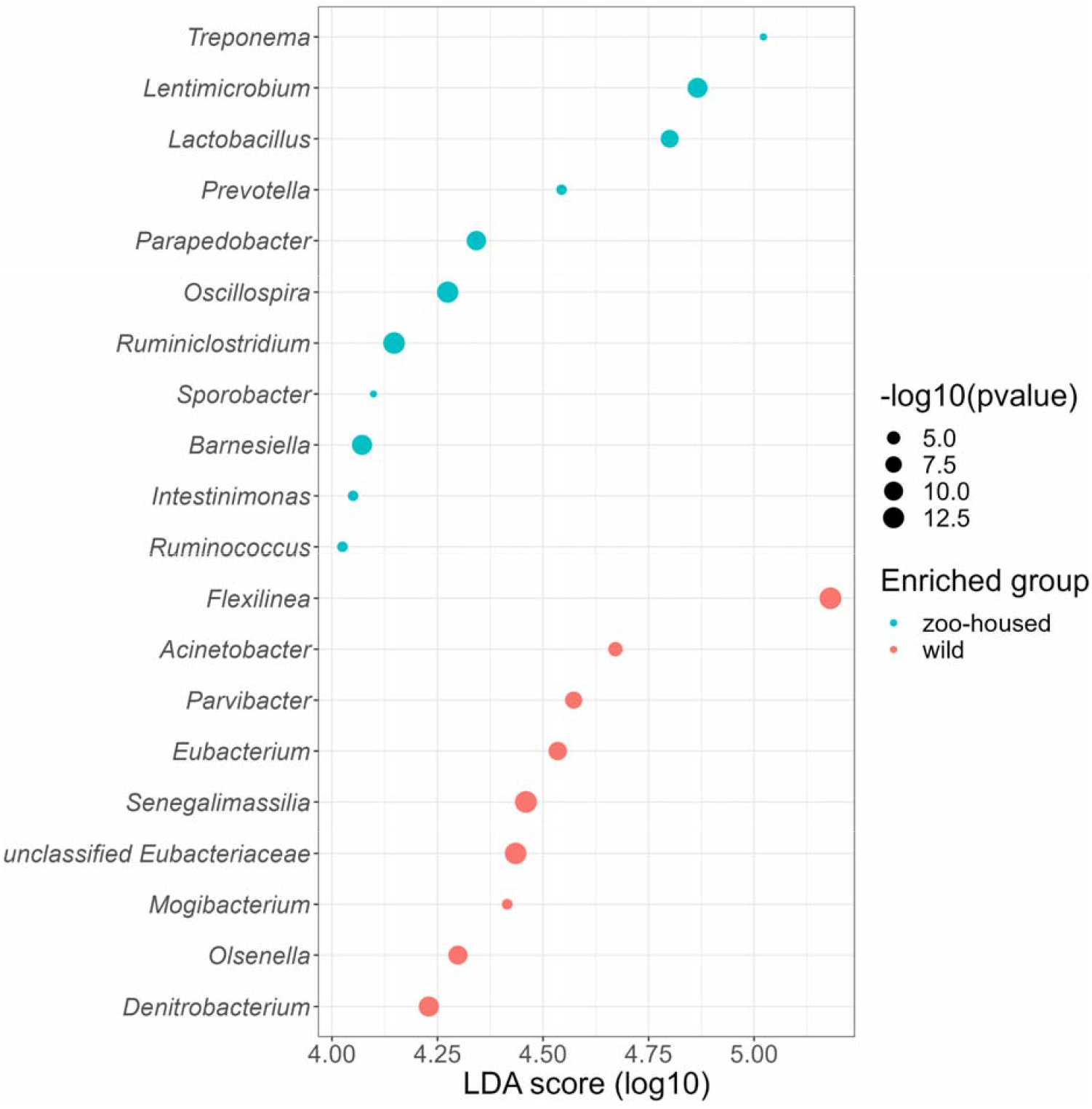
Genera enriched in the microbiota of wild and zoo-housed gorillas. Log10 transformed LDA scores and the corresponding log10 transformed p-values resulting from LefSE analysis at the genus level between WG and ZHG samples. Genera with an LDA score > 4.0 and p-value < 0.05 are shown.

### Altered composition and substantial fungal and archaeal activity in the zoo-housed gorilla gut microbiota

We proceeded with metagenomics (MG) and metatranscriptomics (MT) analysis of four new samples from two time points of the dominant male of the ARTIS gorillas. We used Kraken/Braken2 to determine the taxonomic origin of metagenomic and metatranscriptomic reads. Bacteria accounted for 98.9% of the gorilla gut metagenome. Eukarya (0.07%), Archaea (0.4%), and viruses (0.01%) together accounted for approximately 1% of the remaining genetic material in the gorilla gut. Interestingly, in the gut metatranscriptome only 60.1% of transcripts were of bacterial origin. Eukaryotic transcripts constituted 20.1% of the metatranscriptome, and 19.3% of the transcripts originated from Archaea. The remaining 0.03% of the transcripts were attributed to viruses in MT samples. Taxonomic assignment of the MG and MT samples was in accordance with the 16S rRNA gene amplicon sequencing results of the ARTIS population. Firmicutes and Bacteroides dominate the bacterial microbiota. The major bacterial community composition shifts observed in the 16S rRNA gene amplicon sequencing results were confirmed by the MG and MT reads. Chloroflexi contributed only 0.13% of the MG reads and 0.03% of the MT reads according to the Kraken2/Bracken taxonomy assignment, whereas the high relative abundance of *Lactobacillus* was also reflected in the MT and MG samples:10.5% of the MG reads were assigned to *Lactobacillus*, which accounted for 14.6% of the MT reads.

Almost all eukaryotic transcripts were assigned to the genera *Piromyces, Pecoramyces, and Neocallimastix* (Supplemental Table S4), and all three belonged to Neocallimastigomycetes, a class of anaerobic gut fungi previously identified in the rumen and gut of other herbivores (95% of eukaryotic RNA reads, 19.0% of the total metatranscriptome, and 0.07% of the total metagenome). The remaining 5% of eukaryotic transcripts (1.1% of the total metatranscriptome) are found in soil or plants, such as the pathogenic genus *Melamspora*, which infects *Salix* species (willow tree) that are consumed by zoo-housed gorillas. Thus, these are not considered true members of the gut microbiota, but most likely originated from the passage of food through the gastrointestinal tract. Sequence reads matching the Archaea were present at rather low abundance in the metagenome (0.4 %) but showed much higher abundance in the metatranscriptome (19.3%). They originate from methylotrophic methanogens (genera ‘*Candidatus* Methanoplasma’, 6.8, ‘*Candidatus* Methanomethylophilus’, 3.8%, *Methanomassiliicoccus*, 0.4%), and the more widespread hydrogenotrophic methanogens (genus *Methanobrevibacter*, 8.3%), as indicated in Supplemental Table S5. Metatranscriptome analysis revealed a substantial contribution of anaerobic fungi and archaea to the activity of the gut microbiome, accounting for 38.3% of the assignable metatranscriptome.

### The gorilla microbiome harbors an extensive repertoire of carbohydrate active enzymes

Shifting our perspective from a taxonomic to a functional one, we characterized the carbohydrate active enzyme repertoire of the ZHG gut microbiome. We utilized the classification of the Carbohydrate Active enzyme database (CAZyDB), which provides a sequence-based family classification linking the sequence to the specificity and 3D structure of the enzymes that assemble, modify, and break down oligo- and polysaccharides (Lombard et al., 2014). We identified genes and transcripts belonging to 268 CAZyme families. CAZyDB classifies enzymes into six superfamilies (Supplemental Table S6). Most CAZymes identified in the gorilla gut metagenome (MG) were glycoside hydrolases (GH, 53.7%). The percentual contribution of each GH family to the total of GH families in the CAZy metagenome and metatranscriptome is shown in

Figure 5. The percentages mentioned in the text below represent the percentual contribution of each family to the total CAZyome. Cellulases and hemicellulases were not found among the abundant GH families in the metagenome, accounting for more than 2% of the metagenome. GH13 (4.6%) is involved in starch breakdown. GH2 (3.2%), GH3 (3.2%), GH20 (1.9%), GH1 (1.8%), GH29 (1.6%), GH43 (1.5%), GH92 (1.5%), and GH97 (1.3%) contain oligosaccharide-degrading enzymes. GH78 (1.7%) and GH23 (1.7%) are debranching enzymes. However, in the metatranscriptome, cellulase and hemicellulases were observed within the abundant GH families (Figure 5). We refer to ‘abundant’ as more than 2% of all GH families in the metatranscriptome. Families GH5 (3.5% MT, 0.98% MG), GH9 (2.1% MT, 0.3% MG), GH48 (2.1% MG, 0.01% MT) and GH6 (1.7% MT, not detected in MG) contain cellulases. GH11 (2.6% MG, 0.01% MT) and GH10 (1.4% MT, 0.3% MG) contain endo-hemicellulases. Abundant GH families in the metatranscriptome that are involved in the degradation of oligosaccharides are GH1 (4.6% MT, 1.8% MG) and GH18 (1.9% MT, 0.5% MG). GH57 (1.1% MT, 0.5% MG). GH77 (3.1% MT, 0.9% MG) contains debranching enzymes.

**Figure 5.**
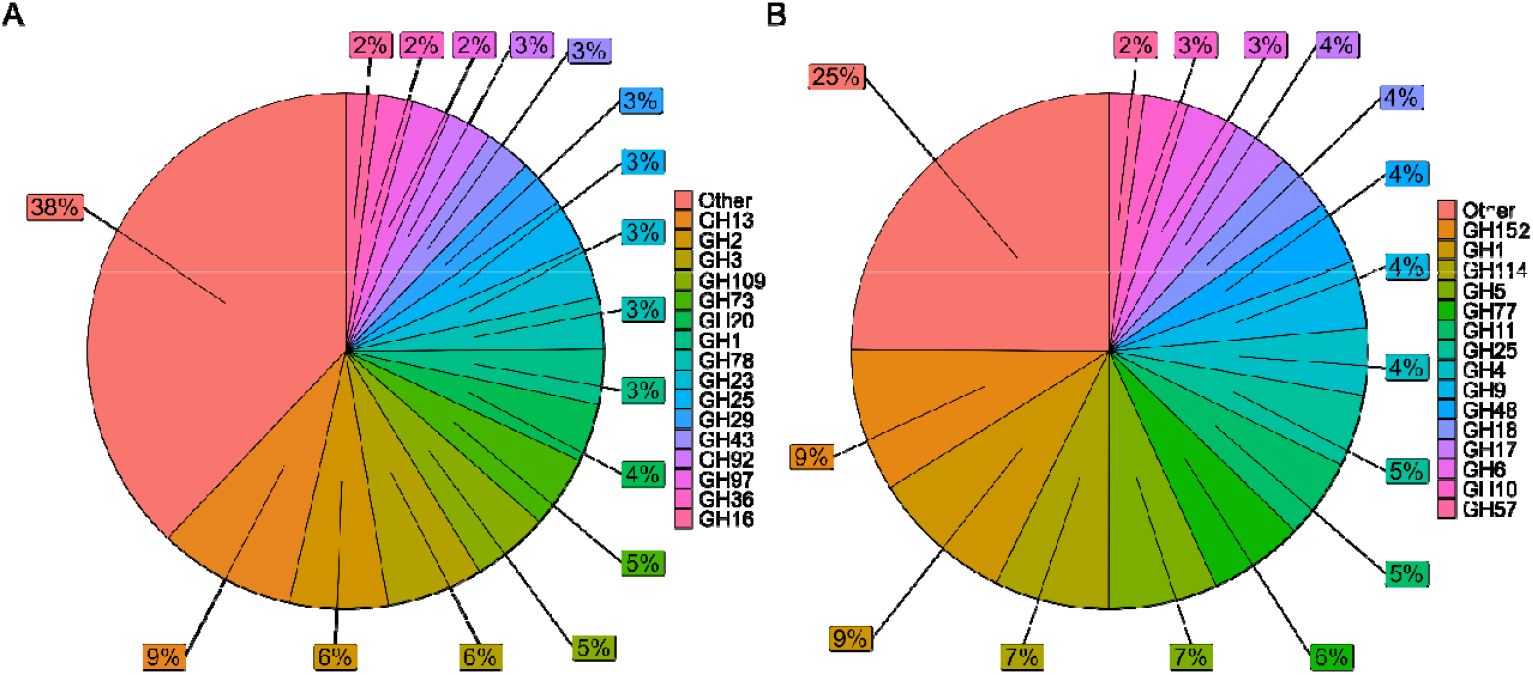
Glycoside hydrolyase families in the gut metagenome and metatranscriptome of zoo-housed gorillas. Percentages represent the percentual contribution of each glycoside hydrolase (GH) family to the total number of GH families found by A) shotgun metagenomics (DNA) and B) metatranscriptomics or RNA-seq (mRNA) on fecal samples of a ZHG.

Mucin degradation by gut microbes requires the expression of GH33 (sialidases) to access mucin glycans (Glover et al., 2022). A total of 0.84% of the metagenomic CAZyome is accounted for by GH33, but no transcripts were observed. Thus, we observed no direct producers of mucin-degrading activity in the ZHG gut microbiome at the time of sampling. Other GH families known to be involved in mucin degradation are GH16, GH29, GH95, GH20, GH2, GH35, GH42, GH98, GH101, GH129, GH89, GH85, and GH84 (Glover et al., 2022); however, these are involved in the general degradation of oligosaccharides, and their presence cannot be considered direct evidence for mucin-degrading activity. Glycosyltransferases (GT) contribute 32.0% of the MG CAZyome and 20.8% of the MT CAZYome (but are not considered further here, since they are involved in polysaccharide synthesis instead of their breakdown). Carbohydrate esterase (CE) families aid in the deacetylation of plant polysaccharides (9.9% MT, 11.5% MG; Supplemental Figure S11). Carbohydrate-binding modules (CBM) target their substrates (mostly cellulose and hemicellulose) and promote prolonged interaction, thereby potentiating the enzymatic activities of other CAZymes. CBMs contributed 3.7% to MG CAZyome and 10.3% to MT CAZYome (Supplemental Figure S12).

Notably, the MT CAZyome did not predict the activity of CAZymes in the MG samples. No (hemi)cellulases were found among the abundant GH families in the MG samples, but the metatranscriptome indicated that they are among the most active families of the CAZyome. The most represented GH family in the metagenome, GH13 (starch degrading), contributed only 0.96% of the total MT CAZyome. Based on metagenome analysis alone, we would have underestimated the (hemi)cellulase activity of the zoo-housed microbiome, just as we would have underestimated the contribution of anaerobic fungi and archaea. The presence and activity of debranching enzymes and oligosaccharide-active enzymes allow for further consumption of degraded plant material by hemicellulases, or breakdown of oligosaccharides directly available in the diet. Interestingly, for families such as GH6 or GH48, none or extremely little MG reads were found, but significant levels of MT reads were detected. We further investigated two of the cellulase families for which this is the case, GH6 and GH48, with ShortBRED, a system for profiling protein families of interest at very high specificity in meta-omic sequencing data (Kaminski et al., 2015).

### The cellulase families GH48 and GH6 originate from anaerobic gut fungi

The GH48 family contains cellulases, and its transcripts were detected in very low quantities in the metagenome. GH6 was found in MT samples but not in MG samples. From the 1405 known GH48 DNA sequences, 110 corresponding protein sequences were deposited in UniProt, of which eight were detected using ShortBRED in our MT data (Supplemental Table S7) and none in the MG data. Within the GH48 family, we detected two transcripts encoding proteins of presumed fungal origin. The first of these transcripts resembles H2BPU6, a putative cellulase of *Neocallimastix patriciarum* that was previously detected in the rumen and functionally described by transcriptomic analysis (Li et al., 2014). The latter maps to Q8J1E3, a Cellulase Cel48A protein known from *Piromyces* sp. strain E2 isolated from the gut of an Indian elephant. Cellulase 48 is the major cellulosome component of *Piromyces* (Steenbakkers et al., 2002). Both species are members of the class Neocallimastigomycetes, which were previously shown to account for 19.0% of taxonomically assignable transcripts. The other GH48 transcripts found in ShortBRED were ascribed to an uncultured actinobacterium from the soil and uncultured bacteria from the rumen. Cellobiohydrolase from GH6 was first identified in *Piromyces rhizinflatus*.

### Hydrogenosomal activity and symbiosis with methanogens

Neocallimastigomycetes possess hydrogenosomes instead of mitochondria (Czajkowski et al., 2020). The malic enzyme (ME) is considered a marker of hydrogenosomal activity (Mentel et al., 2008). Of the 16 malic enzyme proteins attributed to Neocallimastigomycetes in UniProt, transcripts from the ME of *Neocallimastix californiae* and *Piromyces sp*. ((A0A1Y2CXE5 in all samples and A0A1Y3NKG2 in three out of four samples). Other proteins involved in hydrogenosomal metabolism are succinate—CoA ligase subunit alpha (Q7Z941, originating from *Neocallimastix patriciarum*, with the beta subunit being detected in two out of four samples from another genome (P53587, *Neocallimastix frontalis*). Also expected to be in the hydrogenosome are homoacotinase, Aconitate hydratase, NADH dehydrogenase [ubiquinone] flavoprotein 1, Arg5,6 arginine biosynthetic enzyme, Alanine-tRNA ligase, Dynamin-type G domain-containing protein and MSF1-domain-containing protein. An overview of the results is provided in Supplemental Table S8.

From the scientific literature it is known that the hydrogen produced in the hydrogenosome of anaerobic gut fungi is converted to methane by methanogenic archaea (Li et al., 2021). Methyl-coenzyme M reductase (MCR) catalyzes the final step in methanogenesis and is considered a marker gene for methanogenesis (Chen et al., 2020). The alpha, beta, gamma, C, and D subunits of MCR were annotated as COG ids (COG4058, COG4054, COG4057, COG4056, and COG4055, respectively). CPM values of the collective of these COG IDs were 14 to 145 times higher in the metatranscriptome than in the corresponding metagenome samples, indicating an overrepresentation of methanogenic activity in MT samples, in line with the Kraken2/Bracken taxonomy assignment. For methanogenesis, the abundance of genes (metabolic capacity) in the metagenome does not reflect its contribution to transcriptional activity. Our data indicated that anaerobic fungi and their symbiotic methanogens contribute substantially to enzymatic activity in the zoo-housed gut microbiome of gorillas (Supplemental Tables S4 and S5).

## Discussion

In this study, we determined the gut microbiota composition of a population of zoo-housed Western lowland gorillas using 16S rRNA gene V3-V4 amplicon sequencing. Beta-diversity analysis showed that the zoo-housed gorilla microbiota had a significantly distinct composition compared to the wild gorilla microbiota. Their gut microbiota composition showed increased alpha diversity in terms of species richness compared to previously obtained data from wild western lowland gorillas, as indicated by the significantly lower Chao1 estimator, ACE, and Fisher diversity index values. The increased alpha diversity of the ZHG microbiome may be explained by the fact that all the wild gorilla samples were collected during periods of low dietary diversity. As observed in previous studies on primate zoo-housed microbiota, we found that the characteristic members of the gut microbiota of native gorillas vanished. Nine bacterial genera were significantly reduced in the zoo-housed gorillas. The association of some of these genera with the consumption of high-fiber foods (*Flexilinea* and *Olsenella*) consumed during periods of low dietary diversity may explain this (Davenport et al., 2014; Hicks et al., 2018).

A striking observation was the relatively high abundance of *Lactobacillus* species, known to colonize the mammalian gut, in the microbiota of zoo-housed gorillas, particularly in those from ARTIS. An anatomical explanation may be given here: gorillas share a gut anatomy that is similar to that of other monogastric herbivores. A typical type of gut anatomy is a simple stomach, in which the proximal part consists of non-glandular squamous cell epithelium. In horses, it has been shown that *Lactobacillus* species can colonize this area of the stomach and form biofilm-like structures (Yuki et al., 2000). Although this phenomenon has not been studied or observed previously in gorillas, based on the similarity in anatomy of the gorilla and stomach of monogastric herbivores, we hypothesize that the stomach of gorillas may be susceptible to colonization by *Lactobacillus* species as well. Colonization of the non-glandular part of the stomach is believed to be beneficial to the host because it prevents colonization by pathogenic microorganisms (Yuki et al. 2000). This hypothesis of *Lactobacillus* colonization of the gorilla stomach would clarify the presence of these *Lactobacillus* species in our compositional data, but does not explain why the relative abundances are so high compared to the compositional data of wild gorillas. Again, diet most likely explains this observation. While the wild gorilla diet almost exclusively consists of high-fiber fruits and plant material, the zoo-housed gorilla diet inevitably contains high levels of oligosaccharides (Gänzle et al., 2012). They are mainly present in the upper part of the mammalian intestinal tract (Gänzle et al., 2012). The combined effect of easy adhesion to the upper intestinal epithelium and the high availability of oligosaccharides in this part of the gastrointestinal tract may explain why *Lactobacillus* species can flourish in the intestinal tract of zoo-house gorillas. However, based on the relative abundance alone, we cannot draw confident conclusions regarding the absolute abundance and distribution of bacteria in the gastrointestinal tract.

Annotation of metagenomic and metatranscriptomic CAZyome led to the identification of 268 distinct Carbohydrate Active enzyme (CAZyme) families. None of the GH families abundant in the metagenome contained cellulases or hemicellulases. Instead, starch or other oligosaccharide-degrading and debranching enzymes are the most represented GH families in the metagenome. However, the metatranscriptome reveals the activity of multiple cellulases and hemicellulases, next to these oligosaccharide-degrading enzymes and debranching enzymes. From a methodological point of view, it is important to note that we only resorted to metagenomics for functional profiling and concluded that the ZHG microbiota hardly displayed potential for cellulose and hemicellulose degradation. However, metatranscriptomics revealed the activity of cellulases and hemicellulases, and that of Neocallimastigomycetes and methanogens in the gut microbiota of gorillas, which we would not have identified or largely underestimated by metagenomic analysis alone. A first possible explanation for this difference is that, due to the short half-life of mRNA, transcripts active in the upper intestinal tract may be less represented in the metatranscriptome than in the hindgut. Oligosaccharides are mainly digested in the upper intestinal tract, which may explain the overrepresentation of oligosaccharide-degrading enzymes in the metagenome compared with the metatranscriptome. A second explanation could be that some of the (hemi)cellulases are of eukaryotic origin. Interestingly, the activity of at least two families of cellulases (GH6 and GH48) can be attributed to members of the Neocallimastigomycetes, which are underrepresented (0.07%) in the metagenome but contribute 19.0% to the total metatranscriptome. Splicing effects may explain the lack of detection of eukaryotic genes in the metagenome, since methods for metagenomic analysis are tailored to the bacterial microbiome and thus do not account for introns and exons in assembly and contig annotation or lack reference genomes from eukaryotes in their databases (Lind & Pollard, 2021). We presume that this is the reason why no shotgun metagenomics studies have found or mentioned the presence of Neocallimastigomycetes in the gorilla microbiome (Campbell et al., 2020; Hicks et al., 2018).

In fact, only one previous study utilizing ITS1 sequencing has discussed the presence of Neocallimastigomycetes in the gorilla gut (Schulz et al., 2018). The authors of this study recognized that anaerobic fungi are well-known members of the ruminant intestinal tract and are essential for the breakdown of plant material (Czajkowski et al., 2020; Gruninger et al., 2014). ITS1 sequencing results suggested that these fungi were indeed members of the gorilla gut microbiota (Schulz et al., 2018). Later studies utilized 16S rRNA gene amplicon sequencing (which does not detect eukaryotes) or shotgun metagenomics, and did not identify these anaerobic gut fungi. Similarly, in our shotgun metagenomics data, all analyses except for the taxonomic assignment with Kraken2/Bracken did not detect these anaerobic fungi at all. However, our RNA-seq results show that they are highly active in the gorilla gut and that they contribute to cellulase activity. Fungal isolates from the rumen microbiome degraded approximately 70% of dried leaf blades and stems in *in vitro* digestion studies (Akin et al., 1990). Thus, Neocallimastigomycetes may be major players in the degradation of plant cell wall material by the gorilla microbiome and are preserved in the zoo-housed microbiome.

The isolation of Neocallimastigomycetes and novel cellulases from the gorilla gut microbiome could be of great biotechnological interest for biomass degradation. Their polysaccharide-degrading efficiency is the highest in symbiosis with methanogenic archaea (Bauchop & Mountfort, 1981; Li et al., 2021), which is also overrepresented in the metatranscriptome compared to the metagenome. In human microbiome data, metagenome analysis has been found to underestimate methanogenesis when compared to metatranscriptome data (Franzosa et al., 2014, 2018). It should be noted that not all methanogenic activity found in our data originated from methanogens that are symbionts to anaerobic fungi, as these methanogens can also be symbionts to hydrogen-producing bacteria in the gut microbiota (Ma et al., 2020). Based on our analyses, we could not quantify bacterial and fungal contributions to hydrogen production. However, it has been found that anaerobic fungi and methanogens together degrade polysaccharides at a higher rate than a consortium of bacteria and methanogens (Ma et al., 2020), highlighting the biotechnological potential of Neocallimastigomycetes.

The methanogenic activity identified in our study in the gut microbiome of zoo-housed gorillas originated from hydrotrophic and methylotrophic methanogens. We identified three genera of Thermoplasmata as the origin of methylotrophic methanogenesis and the genus *Methanobrevibacter* as the origin of hydrogenotrophic methanogenesis. Because methanogenesis from carbon dioxide consumes four molecules of hydrogen per molecule of methane and methanogenesis from methanol requires only one molecule, methyl-reducing methanogens should have an energetic advantage over hydrogenotrophic methanogens at low hydrogen partial pressures (Vanwonterghem et al., 2016). It remains to be investigated whether the relatively high percentage of methylotrophic methanogens is a specific characteristic of zoo-housed gorillas and whether this is linked to a lower hydrogen pressure.

Our study shows how metatranscriptomics can uncover the contribution of eukaryotes and archaea to the herbivorous gut microbiome. It should be noted that we carried out metagenome and metatranscriptome analysis of four fecal samples from only one zoo-housed gorilla. Ideally, future research would include metatranscriptomic data of wild gorilla microbiomes, although we recognize that this is challenging because fecal samples must be collected as soon as possible after defecation due to the short lifetime of RNA.

In addition, shotgun metagenomic analysis workflows should be improved to detect eukaryotic and prokaryotic genes. In particular, the underrepresentation of gorilla-specific microbes in databases, including the HUMAnN3 database used in this study, poses limitations to the number of unassignable reads. One solution is to map all unassignable reads using BLAST analysis. However, this is computationally very intense if performed for millions of sequences. The prediction of eukaryotic and prokaryotic genes requires a different approach (*i*.*e*., to detect splicing sites in eukaryotic genes), which starts by correctly classifying metagenomic contigs. While some promising tools are being developed to address this, for instance, Whokaryote (Pronk and Medema, 2022), these need to be integrated into larger analysis frameworks.

In conclusion, the zoo-housed microbiome is altered in terms of diversity, composition, and expected function. However, hemi(cellulases), other carbohydrate-active enzymes targeting polysaccharides, and enzymes originating from Neocallimastigomycetes remain active in the zoo-housed microbiome. The abundance of fiber-degrading Chloroflexi was strongly reduced in the microbiota of zoo-housed gorillas; however, the herbivore-characteristic activity of the microbiome was conserved. These results are promising considering conservation efforts and require further investigation to ‘rewild’ the zoo-housed microbiome. Our study highlights the contribution of eukaryotes and archaea to the gorilla gut microbiome, thereby showing the added value of RNA sequencing and the importance of further improvements in metagenomics analysis pipelines for the detection of eukaryotic genes in shotgun metagenomics data.

## Funding

This research project was conducted with internal funding from the ARTIS Heritage, Education, and Academy team.

## Conflict of Interest

The authors declare no conflicts of interest.

## Acknowledgements

The authors would like to thank all gorilla caretakers at ARTIS Amsterdam Royal Zoo for their cooperation and collection of fecal samples. The authors gratefully acknowledge Yaro Laenen and Jasper Buikx from ARTIS-Micropia, and Eline Klaassens, Radhika Bongoni, Mirna Baak, Will Harley from BaseClear for their support to this research project, and Karline Janmaat (ARTIS Chair holder Cognitive Behavioural Ecology at Leiden University) for fruitful discussions.

## Author contributions

Conceptualization, I.H., and R.K.; Methodology and data analysis, I.H., M.v.Z.L., M.B., W.P., R.K; Experiments: M.v.Z.L., R.K.; Data curation, I.H.; Writing -original draft preparation, I.H.; Writing - review and editing, I.H., W.P, R.K.; Supervision, R.K. All authors have read and agreed to the published version of the manuscript.

## Notes

### Competing Interest Statement

The authors have declared no competing interest.

### Summary of Updates

We found two more recent studies by Campbell et al. (2020) and Narat et al. (2020), who have sequenced the gut microbiota of 21 and 43 gorillas, respectively. The datasets include wild and zoo-housed western lowland gorillas. We have integrated these samples into our analysis and removed the comparison with the 454 dataset of Gomez et al. (2015), which was the sole dataset available at the start of our project. To our knowledge, our revised work comprises the largest sample amount available: a total 79 samples (including 15 samples from our study) to date and provides a more solid basis for our conclusions. Furthermore, we have revised Figure 1 (rarefaction curve) and moved Figure 1 to the supplemental data, inserted a PCA plot of all zoo-housed and wild gorilla microbiota compositions (new Figure 1), alpha diversity indices (new Figure 2), heatmap of the top 10 genera of zoo-housed and wild gorillas (new Figure 3), and a figure with enriched genera in the microbiota of wild and zoo-housed gorillas (new Figure 4).

## References

Akin, D. E., Borneman, W. S., & Lyon, C. E. (1990). Degradation of leaf blades and stems by monocentric and polycentric isolates of ruminal fungi. Animal Feed Science and Technology, 31(3–4), 205–221. https://doi.org/10.1016/0377-8401(90)90125-r

Andrews, S. (2010). FastQC: a quality control tool for high throughput sequence data. URL http://www.bioinformatics.babraham.ac.uk/projects/fastqc

Bauchop, T., & Mountfort, D. O. (1981). Cellulose fermentation by a rumen anaerobic fungus in both the absence and the presence of rumen methanogens. Applied and Environmental Microbiology, 42(6), 1103–1110. https://doi.org/10.1128/aem.42.6.1103-1110.1981

Beghini, F., McIver, L. J., Blanco-Míguez, A., Dubois, L., Asnicar, F., Maharjan, S., Mailyan, A., Manghi, P., Scholz, M., Thomas, A. P., Valles-Colomer, M., Weingart, G., Zhang, Y., Zolfo, M., Huttenhower, C., Franzosa, E. A., & Segata, N. (2021). Integrating taxonomic, functional, and strain-level profiling of diverse microbial communities with bioBakery 3. eLife, 10. https://doi.org/10.7554/elife.65088

Campbell, T. P., Sun, X., Patel, V. H., Sanz, C., Morgan, D., & Dantas, G. (2020). The microbiome and resistome of chimpanzees, gorillas, and humans across host lifestyle and geography. The ISME Journal, 14(6), 1584. https://doi.org/10.1038/S41396-020-0634-2

Cao, Y., Dong, Q., Wang, D., Zhang, P., Liu, Y., & Niu, C. (2022). microbiomeMarker: an R/Bioconductor package for microbiome marker identification and visualization. Bioinformatics, 38(16), 4027–4029. https://doi.org/10.1093/bioinformatics/btac438

Chen, H., Gan, Q., & Fan, C. (2020). Methyl-coenzyme M reductase and its post-translational modifications. Frontiers in Microbiology, 11, 578356. https://doi.org/10.3389/fmicb.2020.578356

Chen, S., Zhou, Y., Chen, Y., & Gu, J. (2018). Fastp: An ultra-fast all-in-one FASTQ preprocessor. Bioinformatics, 34(17), i884–i890. https://doi.org/10.1093/bioinformatics/bty560

Clayton, J. B., Gomez, A., Amato, K., Knights, D., Travis, D. A., Blekhman, R., Knight, R., Leigh, S., Stumpf, R., Wolf, T., Glander, K. E., Cabana, F., & Johnson, T. J. (2018). The gut microbiome of nonhuman primates: Lessons in ecology and evolution. American Journal of Primatology, 80(6). https://doi.org/10.1002/ajp.22867

Clayton, J. B., Vangay, P., Huang, H., Ward, T., Hillmann, B. M., Al-Ghalith, G. A., Travis, D. A., Long, H. T., van Tuan, B., van Minh, V., Cabana, F., Nadler, T., Toddes, B., Murphy, T., Glander, K. E., Johnson, T. J., & Knights, D. (2016). Captivity humanizes the primate microbiome. Proceedings of the National Academy of Sciences of the United States of America, 113(37), 10376–10381. https://doi.org/10.1073/pnas.1521835113

Cole, J. R., Wang, Q., Fish, J. A., Chai, B., McGarrell, D. M., Sun, Y., Brown, C. T., Porras-Alfaro, A., Kuske, C. R., & Tiedje, J. M. (2014). Ribosomal Database Project: data and tools for high throughput rRNA analysis. Nucleic Acids Research, 42(Database issue). https://doi.org/10.1093/nar/gkt1244

Czajkowski, R., Elshahed, M. S., Singh, B., Hess, M., Edwards, J. E., Paul, S. S., Puniya, A. K., van der Giezen, M., Shaw, C., & Fliegerová, K. (2020). Anaerobic Fungi: Past, Present, and Future. https://doi.org/10.3389/fmicb.2020.584893

Davenport, E. R., Mizrahi-Man, O., Michelini, K., Barreiro, L. B., Ober, C., & Gilad, Y. (2014). Seasonal Variation in Human Gut Microbiome Composition. PLOS ONE, 9(3), e90731. https://doi.org/10.1371/journal.pone.0090731

di Tommaso, P., Chatzou, M., Floden, E. W., Barja, P. P., Palumbo, E., & Notredame, C. (2017). Nextflow enables reproducible computational workflows. Nature Biotechnology, 35(4), 316–319. https://doi.org/10.1038/nbt.3820

Dixon, P. (2003). VEGAN, a package of R functions for community ecology. Journal of Vegetation Science, 14(6), 927–930. https://doi.org/10.1111/j.1654-1103.2003.tb02228.x

Dobin, A., Davis, C. A., Schlesinger, F., Drenkow, J., Zaleski, C., Jha, S., Batut, P., Chaisson, M., & Gingeras, T. R. (2013). STAR: ultrafast universal RNA-seq aligner. Bioinformatics (Oxford, England), 29(1), 15–21. https://doi.org/10.1093/bioinformatics/bts635

Edgar, R. C., & Bateman, A. (2010). Search and clustering orders of magnitude faster than BLAST. Bioinformatics, 26(19), 2460–2461. https://doi.org/10.1093/bioinformatics/btq461

Foster, Z. S. L., Sharpton, T. J., & Grünwald, N. J. (2017). Metacoder: An R package for visualization and manipulation of community taxonomic diversity data. PLoS Computational Biology, 13(2). https://doi.org/10.1371/journal.pcbi.1005404

Franzosa, E. A., McIver, L. J., Rahnavard, G., Thompson, L. R., Schirmer, M., Weingart, G., Lipson, K. S., Knight, R., Caporaso, J. G., Segata, N., & Huttenhower, C. (2018). Species-level functional profiling of metagenomes and metatranscriptomes. Nature Methods, 15(11), 962–968. https://doi.org/10.1038/s41592-018-0176-y

Franzosa, E. A., Morgan, X. C., Segata, N., Waldron, L., Reyes, J., Earl, A. M., Giannoukos, G., Boylan, M. R., Ciulla, D., Gevers, D., Izard, J., Garrett, W. S., Chan, A. T., & Huttenhower, C. (2014). Relating the metatranscriptome and metagenome of the human gut. Proceedings of the National Academy of Sciences of the United States of America, 111(22). https://doi.org/10.1073/pnas.1319284111

Fu, L., Niu, B., Zhu, Z., Wu, S., & Li, W. (2012). CD-HIT: accelerated for clustering the next-generation sequencing data. Bioinformatics, 28(23), 3150. https://doi.org/10.1093/bioinformatics/bts565

Gänzle, M. G., Follador, R., & van Derlinden, E. (2012). Metabolism of oligosaccharides and starch in lactobacilli: a review. Frontiers in microbiology, 3, 340.https://doi.org/10.3389/fmicb.2012.00340

Glover, J. S., Ticer, T. D., & Engevik, M. A. (2022). Characterizing the mucin-degrading capacity of the human gut microbiota. Scientific Reports 2022 12:1, 12(1), 1–14. https://doi.org/10.1038/s41598-022-11819-z

Gomez, A. (2014). The gut microbiome of the western lowland gorilla (Gorilla gorilla gorilla): implications for overall ecology. PhD-thesis. University of Illinois at Urbana-Champaign, Champaign, IL, USA.

Gruninger, R. J., Puniya, A. K., Callaghan, T. M., Edwards, J. E., Youssef, N., Dagar, S. S., Fliegerova, K., Griffith, G. W., Forster, R., Tsang, A., Mcallister, T., & Elshahed, M. S. (2014). Anaerobic fungi (phylum Neocallimastigomycota): advances in understanding their taxonomy, life cycle, ecology, role and biotechnological potential. FEMS Microbiology Ecology, 90(1), 1–17. https://doi.org/10.1111/1574-6941.12383

Hicks, A. L., Lee, K. J., Couto-Rodriguez, M., Patel, J., Sinha, R., Guo, C., Olson, S. H., Seimon, A., Seimon, T. A., Ondzie, A. U., Karesh, W. B., Reed, P., Cameron, K. N., Lipkin, W. I., & Williams, B. L. (2018). Gut microbiomes of wild great apes fluctuate seasonally in response to diet. Nature Communications 2018 9:1, 9(1), 1–18. https://doi.org/10.1038/s41467-018-04204-w

Cantalapiedra, C. P., Hernández-Plaza, A., Letunic, I., Bork, P., & Huerta-Cepas, J. (2021). eggNOG-mapper v2: functional annotation, orthology assignments, and domain prediction at the metagenomic scale. Molecular biology and evolution, 38(12), 5825–5829.

Huerta-Cepas, J., Szklarczyk, D., Heller, D., Hernández-Plaza, A., Forslund, S. K., Cook, H., Mende, D. R., Letunic, I., Rattei, T., Jensen, L. J., von Mering, C., & Bork, P. (2019). eggNOG 5.0: a hierarchical, functionally and phylogenetically annotated orthology resource based on 5090 organisms and 2502 viruses. Nucleic Acids Research, 47(D1), D309–D314. https://doi.org/10.1093/nar/gky1085

Houtkamp, I. (2022). Gorilla, Github repository, https://github.com/imhoutkamp/Gorilla

Kaminski, J., Gibson, M. K., Franzosa, E. A., Segata, N., Dantas, G., & Huttenhower, C. (2015). High-Specificity Targeted Functional Profiling in Microbial Communities with ShortBRED. PLOS Computational Biology, 11(12), e1004557. https://doi.org/10.1371/journal.pcbi.1004557

Klindworth, A., Pruesse, E., Schweer, T., Peplies, J., Quast, C., Horn, M., & Glöckner, F. O. (2013). Evaluation of general 16S ribosomal RNA gene PCR primers for classical and next-generation sequencing-based diversity studies. Nucleic Acids Research, 41(1). https://doi.org/10.1093/nar/gks808

Kopylova, E., Noé, L., & Touzet, H. (2012). SortMeRNA: Fast and accurate filtering of ribosomal RNAs in metatranscriptomic data. Bioinformatics, 28(24), 3211–3217. https://doi.org/10.1093/bioinformatics/bts611

Langmead, B., & Salzberg, S. L. (2012). Fast gapped-read alignment with Bowtie 2. Nature Methods, 9(4), 357–359. https://doi.org/10.1038/nmeth.1923

Li, D., Liu, C. M., Luo, R., Sadakane, K., & Lam, T. W. (2015). MEGAHIT: an ultra-fast single-node solution for large and complex metagenomics assembly via succinct de Bruijn graph. Bioinformatics, 31(10), 1674–1676.https://doi.org/10.1093/bioinformatics/btv033

Li, J., Su, X., Tian, Y., Dong, Z., Hu, S., Huang, L., & Dai, X. (2014). Gene diversity of the bacterial 48 family glycoside hydrolase (GH48) in rumen environment. Wei Sheng Wu Xue Bao = Acta Microbiologica Sinica, 54(1), 53–61. https://europepmc.org/article/med/24783854

Li, Y., Jin, W., Cheng, Y., & Zhu, W. (2016). Effect of the Associated Methanogen Methanobrevibacter thaueri on the Dynamic Profile of End and Intermediate Metabolites of Anaerobic Fungus Piromyces sp. F1. Current Microbiology, 73(3), 434–441. https://doi.org/10.1007/S00284-016-1078-9

Li, Y., Meng, Z., Xu, Y., Shi, Q., Ma, Y., Aung, M., Cheng, Y., & Zhu, W. (2021). Interactions between Anaerobic Fungi and Methanogens in the Rumen and Their Biotechnological Potential in Biogas Production from Lignocellulosic Materials. Microorganisms 2021, Vol. 9, Page 190, 9(1), 190. https://doi.org/10.3390/microorganisms9010190

Lu, J., Breitwieser, F.P., Thielen, P., Salzberg, S.L. (2017). Bracken: estimating species abundance in metagenomics data. PeerJ Computer Science, 3:e104. https://doi.org/10.7717/peerj-cs.104

Lind, A. L., & Pollard, K. S. (2021). Accurate and sensitive detection of microbial eukaryotes from whole metagenome shotgun sequencing. Microbiome, 9(1). https://doi.org/10.1186/s40168-021-01015-y

Lombard, V., Golaconda Ramulu, H., Drula, E., Coutinho, P. M., & Henrissat, B. (2014). The carbohydrate-active enzymes database (CAZy) in 2013. Nucleic acids research, 42(D1), D490–D495. https://doi.org/10.1093/nar/gkt1178

Ma, Y., Li, Y., Li, Y., Cheng, Y., & Zhu, W. (2020). The enrichment of anaerobic fungi and methanogens showed higher lignocellulose degrading and methane producing ability than that of bacteria and methanogens. World Journal of Microbiology and Biotechnology, 36(9). https://doi.org/10.1007/S11274-020-02894-3

Masi, S. (2011). Differences in gorilla nettle-feeding between captivity and the wild: Local traditions, species typical behaviors or merely the result of nutritional deficiencies? Animal Cognition, 14(6), 921–925. https://doi.org/10.1007/s10071-011-0457-7/figures/1

McMurdie, P. J., & Holmes, S. (2013). phyloseq: An R Package for Reproducible Interactive Analysis and Graphics of Microbiome Census Data. PLOS ONE, 8(4), e61217. https://doi.org/10.1371/journal.pone.0061217

Mentel, M., Zimorski, V., Haferkamp, P., Martin, W., & Henze, K. (2008). Protein import into hydrogenosomes of Trichomonas vaginalis involves both N-terminal and internal targeting signals: a case study of thioredoxin reductases. Eukaryotic cell, 7(10), 1750–1757. https://doi.org/10.1128/EC.00206-08

Moeller, A. H., Caro-Quintero, A., Mjungu, D., Georgiev, A. v., Lonsdorf, E. v., Muller, M. N., Pusey, A. E., Peeters, M., Hahn, B. H., & Ochman, H. (2016). Cospeciation of gut microbiota with hominids. Science, 353(6297), 380–382. https://doi.org/10.1126/science.aaf3951

Moeller, A. H., & Sanders, J. G. (2020). Roles of the gut microbiota in the adaptive evolution of mammalian species. Philosophical Transactions of the Royal Society B: Biological Sciences, 375(1808), 20190597. https://doi.org/10.1098/rstb.2019.0597

Narat, V., Amato, K. R., Ranger, N., Salmona, M., Mercier-Delarue, S., Rupp, S., Ambata, P., Njouom, R., Simon, F., Giles-Vernick, T., & LeGoff, J. (2020). A multi-disciplinary comparison of great ape gut microbiota in a Central African Forest and European Zoo. Scientific Reports, 10(1). https://doi.org/10.1038/s41598-020-75847-3

Nishida, A. H., & Ochman, H. (2021). Captivity and the co-diversification of great ape microbiomes. Nature Communications 2021 12:1, 12(1), 1–10. https://doi.org/10.1038/s41467-021-25732-y

Pronk, L. J. U., & Medema, M.H. (2022). Whokaryote: distinguishing eukaryotic and prokaryotic contigs in metagenomes based on gene structure. Microbial Genomics, 8(5), mgen000823. https://doi.org/10.1099/mgen.0.000823.

Quast, C., Pruesse, E., Yilmaz, P., Gerken, J., Schweer, T., Yarza, P., Peplies, J., & Glöckner, F. O. (2013). The SILVA ribosomal RNA gene database project: improved data processing and web-based tools. Nucleic Acids Research, 41(D1), D590–D596. https://doi.org/10.1093/nar/gks1219

Schulz, D., Qablan, M. A., Profousova-Psenkova, I., Vallo, P., Fuh, T., Modry, D., Piel, A. K., Stewart, F., Petrzelkova, K. J., & Fliegerová, K. (2018). Anaerobic Fungi in Gorilla (Gorilla gorilla gorilla) Feces: an Adaptation to a High-Fiber Diet? International Journal of Primatology, 39(4), 567–580. https://doi.org/10.1007/S10764-018-0052-8

Segata, N., Izard, J., Waldron, L., Gevers, D., Miropolsky, L., Garrett, W. S., & Huttenhower, C. (2011). Metagenomic biomarker discovery and explanation. Genome biology, 12, 1–18. https://doi.org/10.1186/gb-2011-12-6-r60

Seemann, T. (2014). Prokka: Rapid prokaryotic genome annotation. Bioinformatics, 30(14), 2068–2069. https://doi.org/10.1093/bioinformatics/btu153

Sonnenburg, E. D., & Sonnenburg, J. L. (2019a). The ancestral and industrialized gut microbiota and implications for human health. Nature Reviews Microbiology 2019 17:6, 17(6), 383–390. https://doi.org/10.1038/s41579-019-0191-8

Sonnenburg, J. L., & Sonnenburg, E. D. (2019b). Vulnerability of the industrialized microbiota. Science, 366(6464). https://doi.org/10.1126/science.aaw9255

Steenbakkers, P. J. M., Freelove, A., van Cranenbroek, B., Sweegers, B. M. C., Harhangi, H. R., Vogels, G. D., Hazlewood, G. P., Gilbert, H. J., & Op Den Camp, H. J. M. (2002). The major component of the cellulosomes of anaerobic fungi from the genus piromyces is a family 48 glycoside hydrolase. Mitochondrial DNA, 13(6), 313–320. https://doi.org/10.1080/1042517021000024191

Sun L, Toyonaga M, Ohashi A, Tourlousse DM, Matsuura N, Meng XY, Tamaki H, Hanada S, Cruz R, Yamaguchi T, & Sekiguchi (2016a). Lentimicrobium saccharophilum gen. nov., sp. nov., a strictly anaerobic bacterium representing a new family in the phylum Bacteroidetes, and proposal of Lentimicrobiaceae fam. nov. International journal of systematic and evolutionary microbiology, 66(7), 2635–2642. https://doi.org/10.1099/ijsem.0.001103

Sun, L., Toyonaga, M., Ohashi, A., Matsuura, N., Tourlousse, D. M., Meng, X. Y., Tamaki H, Hanada S, Cruz R, Yamaguchi T, & Sekiguchi Y. (2016b). Isolation and characterization of Flexilinea flocculi gen. nov., sp. nov., a filamentous, anaerobic bacterium belonging to the class Anaerolineae in the phylum Chloroflexi. International journal of systematic and evolutionary microbiology, 66(2), 988–996. https://doi.org/10.1099/ijsem.0.000822

Wood, D.E., Lu, J., Langmead, B. (2019). Improved metagenomic analysis with Kraken 2. Genome Biology 20(257): 762302. https://doi.org/10.1186/s13059-019-1891-0

Yuki, N., Shimazaki, T., Kushiro, A., Watanabe, K., Uchida, K., Yuyama, T., & Morotomi, M. (2000). Colonization of the Stratified Squamous Epithelium of the Nonsecreting Area of Horse Stomach by Lactobacilli. Applied and Environmental Microbiology, 66(11), 5030. https://doi.org/10.1128/AEM.66.11.5030-5034.2000

Zhang, H., Yohe, T., Huang, L., Entwistle, S., Wu, P., Yang, Z., Busk, P. K., Xu, Y., & Yin, Y. (2018). dbCAN2: a meta server for automated carbohydrate–active enzyme annotation. Nucleic Acids Research, 46(W1), W95–W101. https://doi.org/10.1093/nar/gky418

Zhang, X., Li, L., Butcher, J., Stintzi, A., & Figeys, D. (2019). Advancing functional and translational microbiome research using meta-omics approaches. Microbiome 2019 7:1, 7(1), 1–12. https://doi.org/10.1186/S40168-019-0767-6

