## Supplementary material for "Multiomics characterization of the of the zoo-housed gorilla gut microbiome reveals bacterial community compositions shifts, fungal cellulose-degrading, and archaeal methanogenic activity"

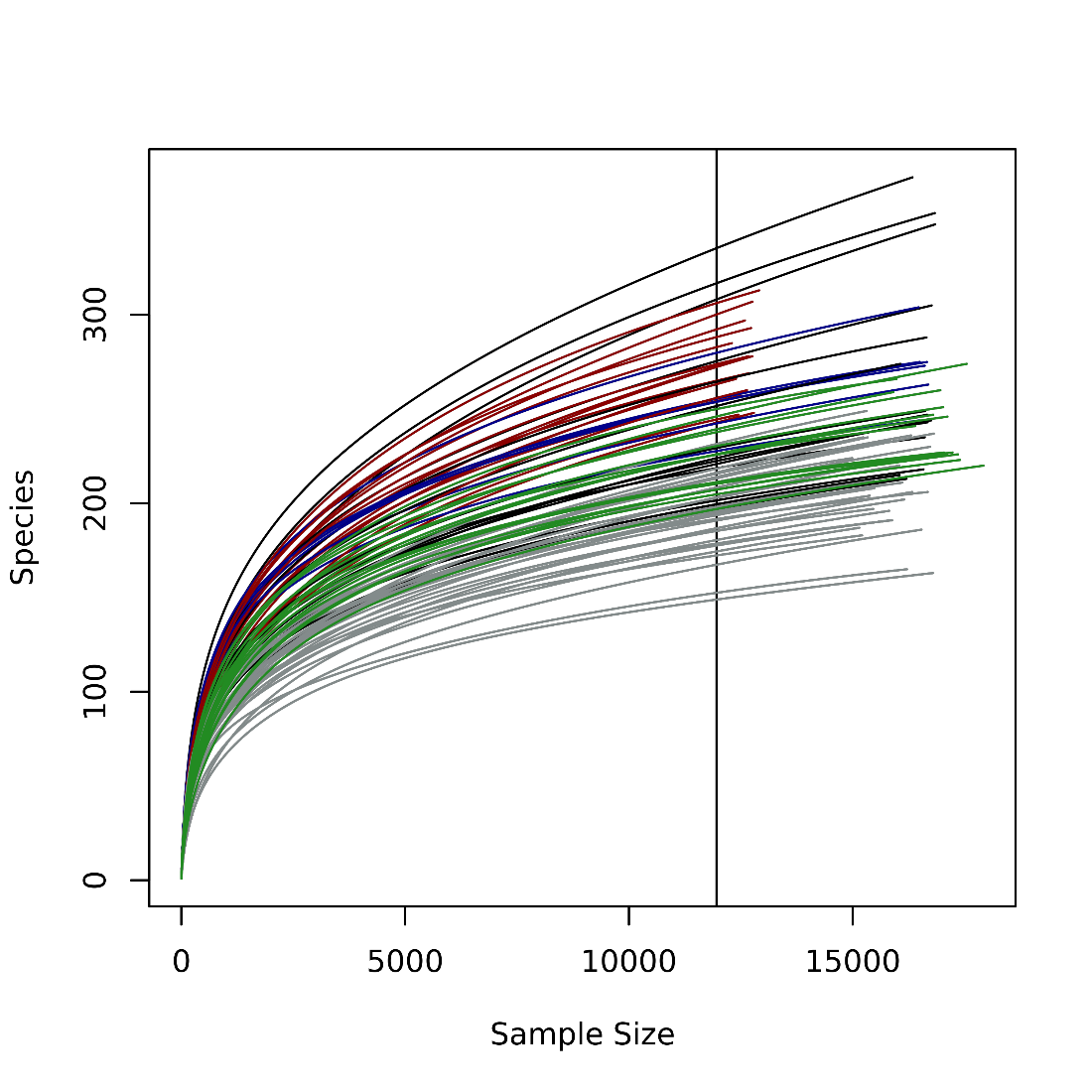

**Figure S1: Rarefaction curve of 16S rRNA gene amplicon sequence data.** Microbiota of zoo-housed gorillas, red (this study); Microbiota of wild gorillas, blue (Campbell et al.,2020); Microbiota of zoo-housed gorillas, green (Campbell *et al.*,2020); Microbiota of wild gorillas, black (Narat *et al.*,2020); Microbiota of zoo-housed gorillas, grey (Narat et al.,2020).

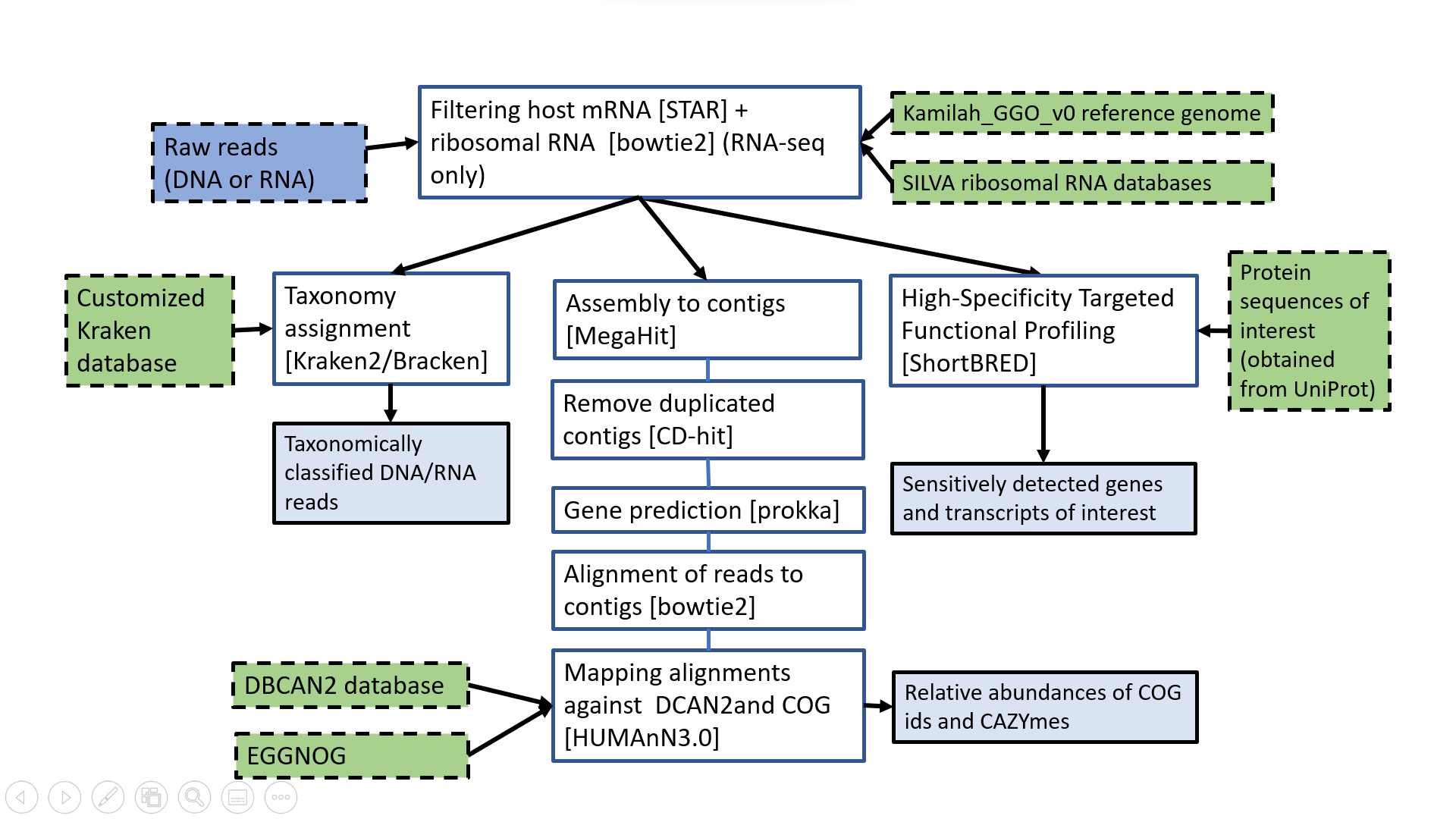

**Figure S2: Shotgun metagenomics/RNA-seq analysis workflow.** The workflow below summarizes the analysis of DNA and RNA reads obtained through shotgun metagenomics and RNA-seq, respectively, as described in detail in the methods section.

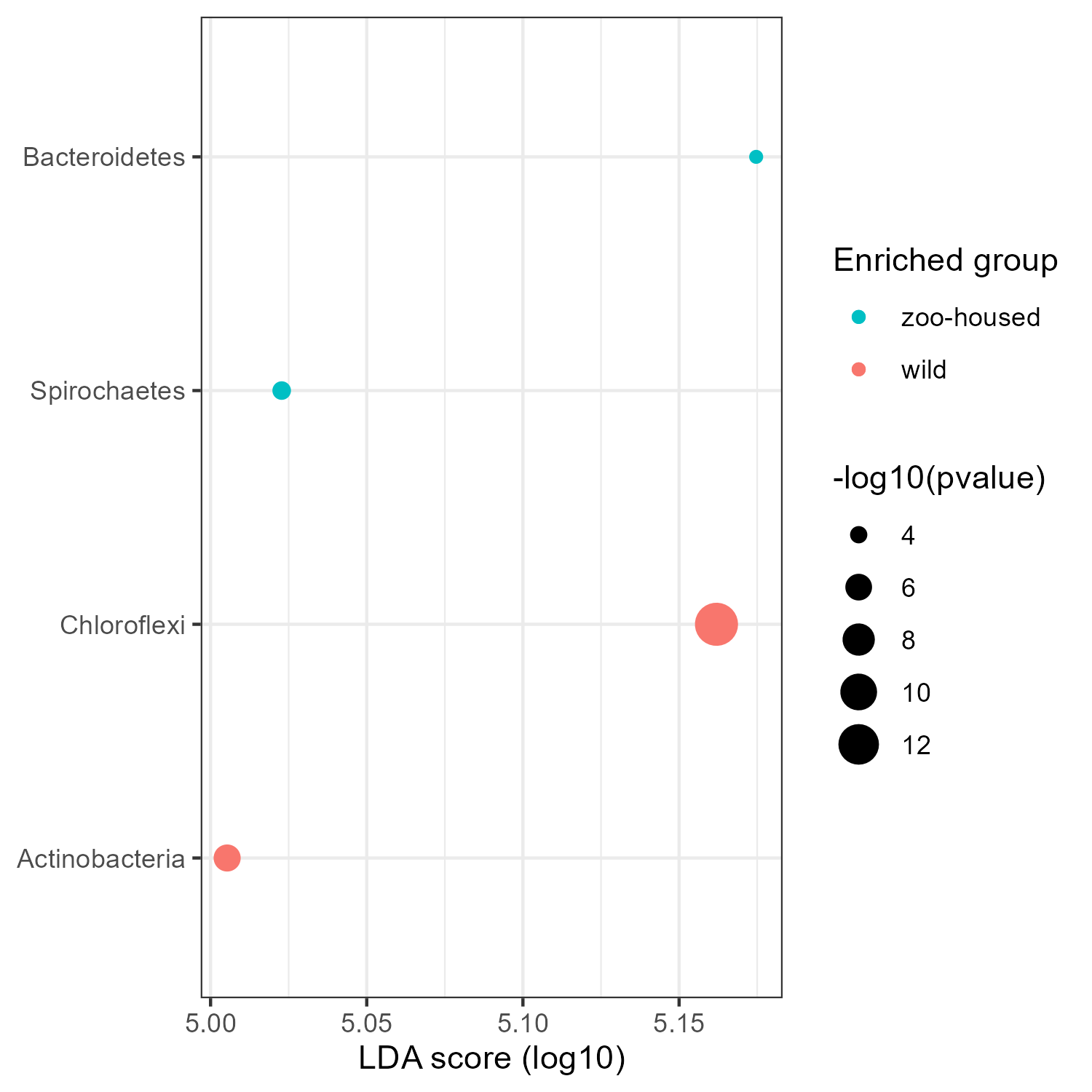

**Figure S3. Phyla identified as significantly enriched in wild or zoo-housed samples by LEfSE analysis.** Log10 transformed LDA scores above 4 and corresponding log10 transformed p-values, resulting from LefSE analysis at genus level between WG and ZHG samples. Genera with an LDA score > 4.0 and p-value < 0.05 are shown.

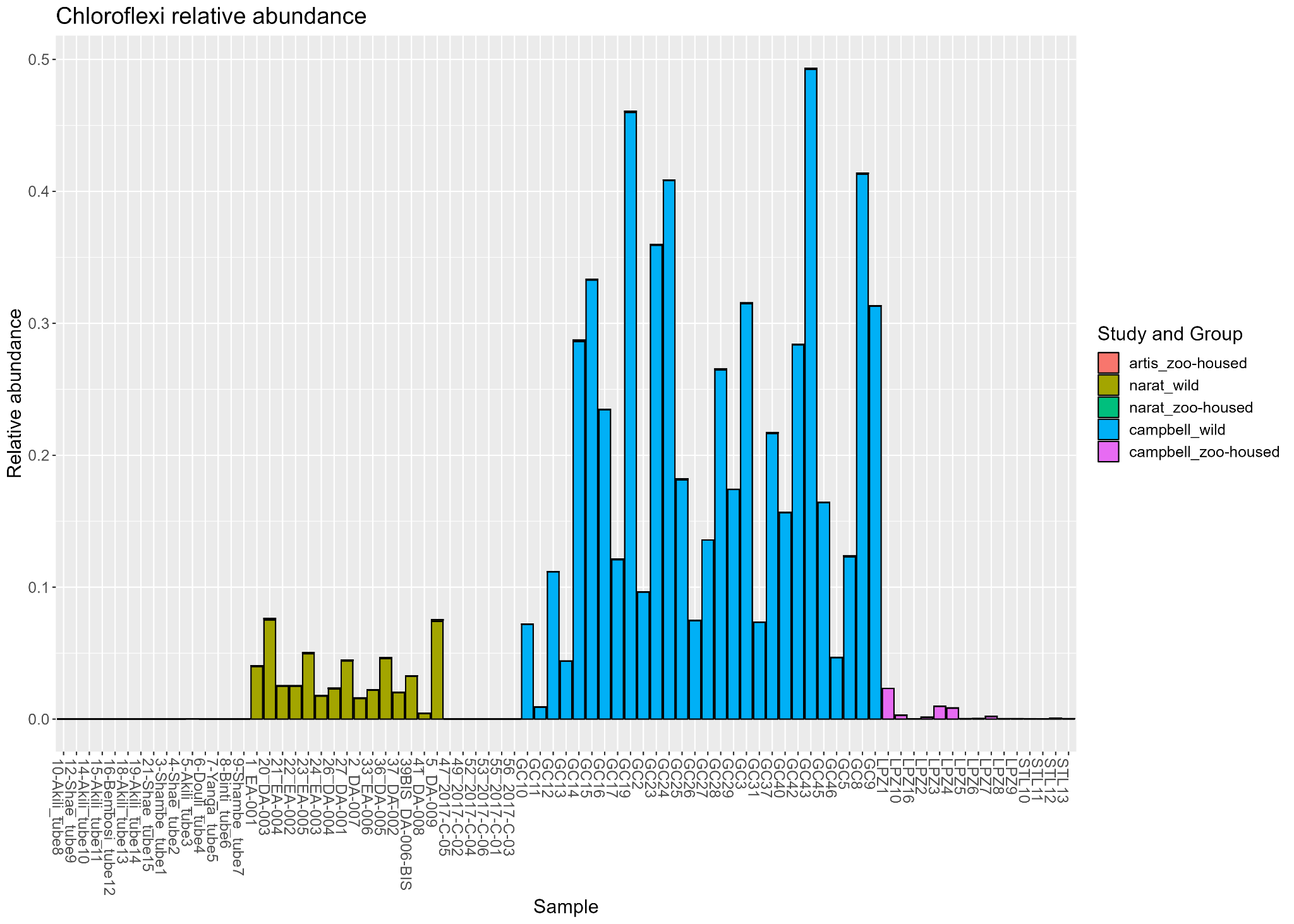

**Figure S4: Relative abundances of the phylum Chloroflexi.** Relative abundances (between 0 and 1) are shown per sample. Colors indicate which study a fecal sample belongs to and whether it originates from a wild or zoo-housed gorilla.

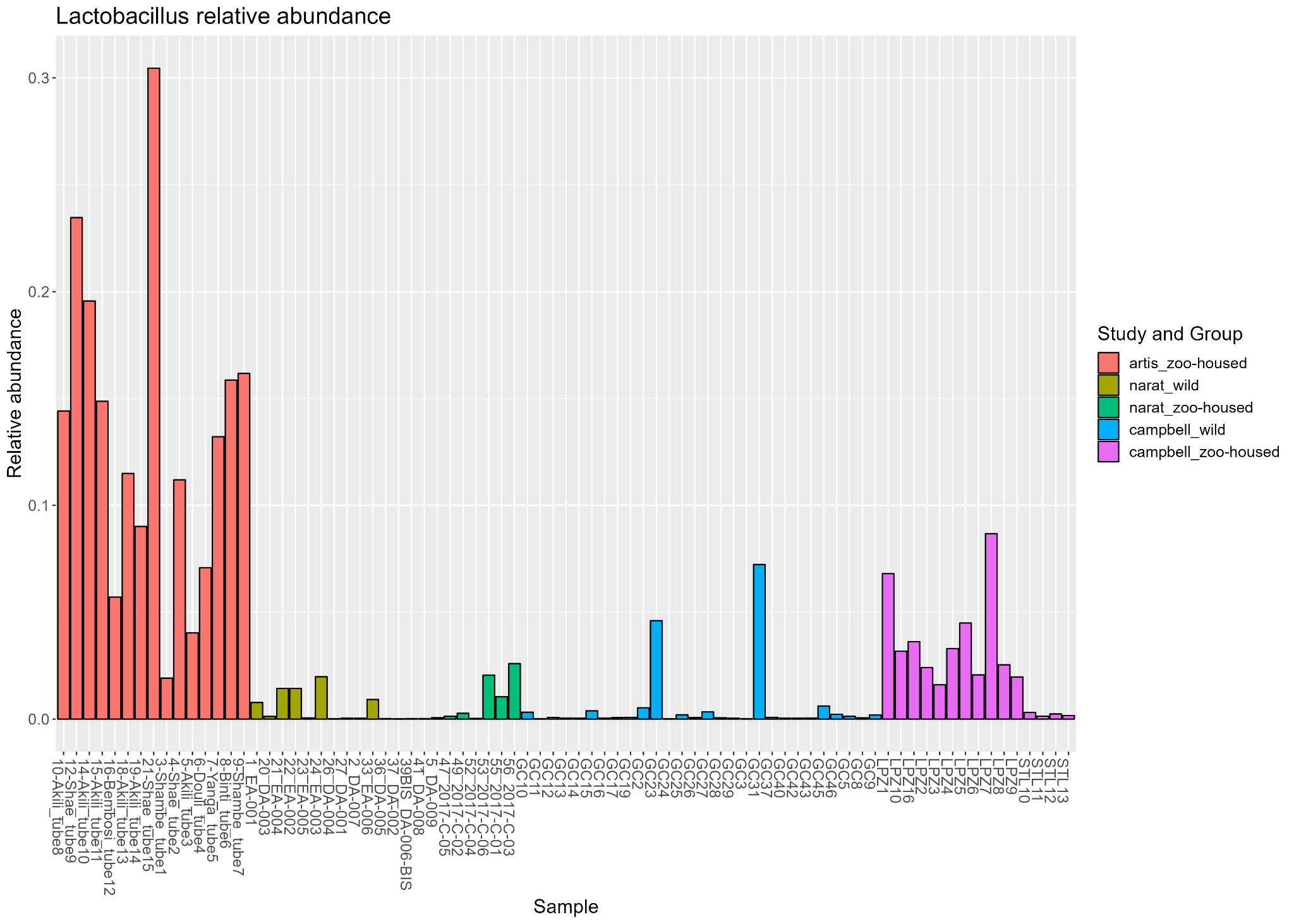

**Figure S5: Relative abundances of the genus *Lactobacillus*.** Relative abundances (between 0 and 1) are shown per sample. Colors indicate which study a fecal sample belongs to and whether it originates from a wild or zoo-housed gorilla.

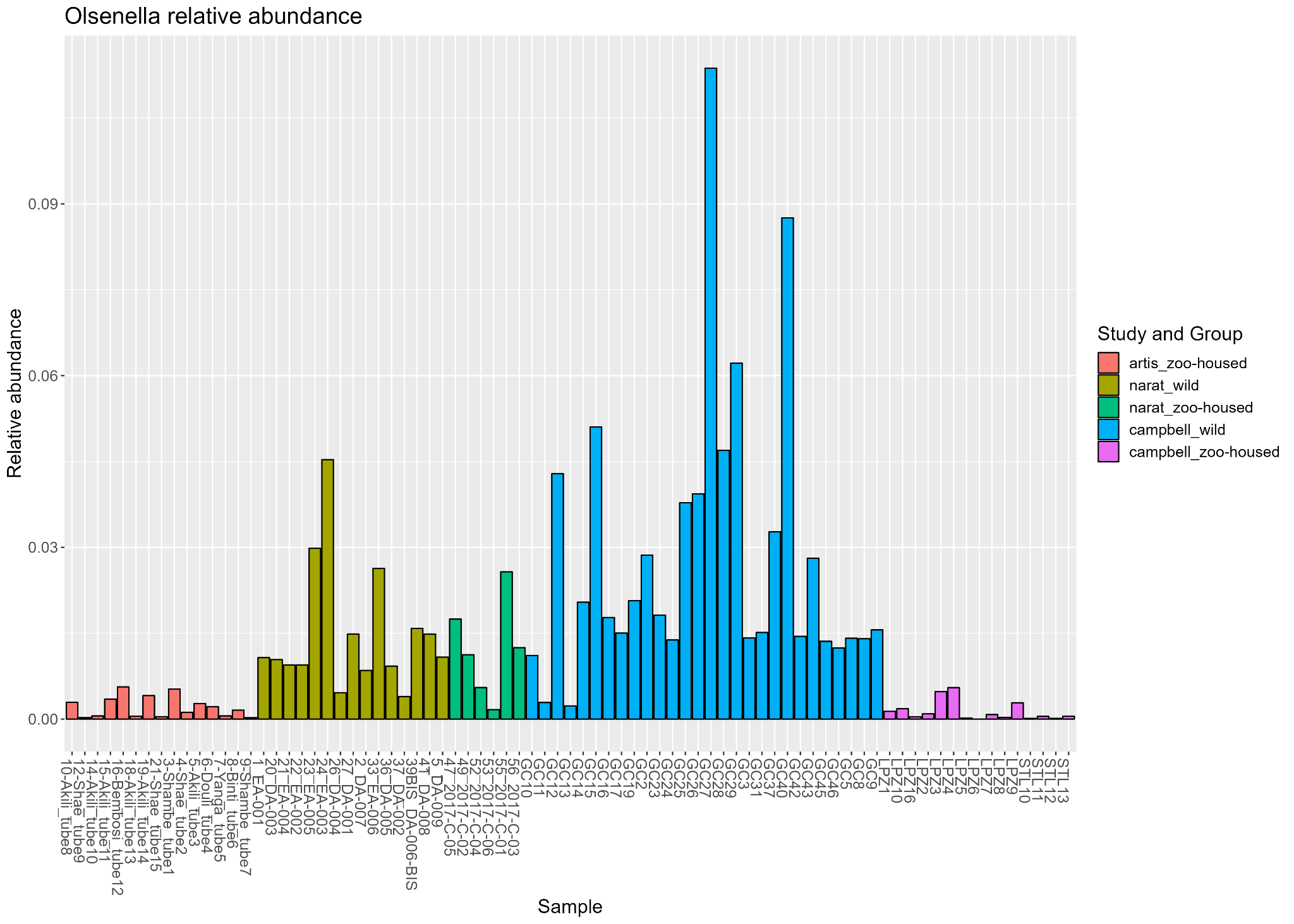

**Figure S6: Relative abundances of the genus *Olsenella*.** Relative abundances (between 0 and 1) are shown per sample. Colors indicate which study a fecal sample belongs to and whether it originates from a wild or zoo-housed gorilla.

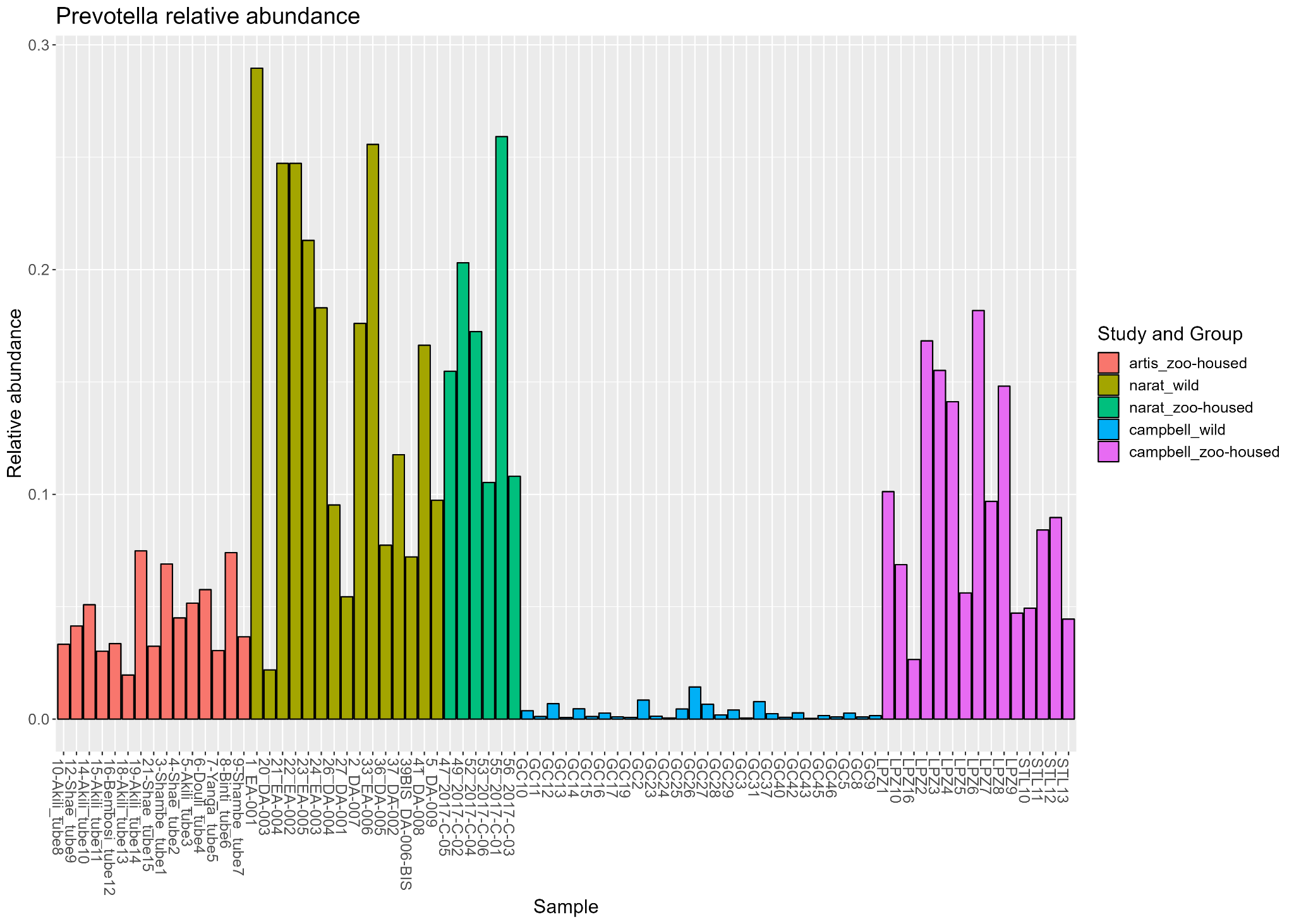

**Figure S7: Relative abundances of the genus *Prevotella*.** Relative abundances (between 0 and 1) are shown per sample. Colors indicate which study a fecal sample belongs to and whether it originates from a wild or zoo-housed gorilla.

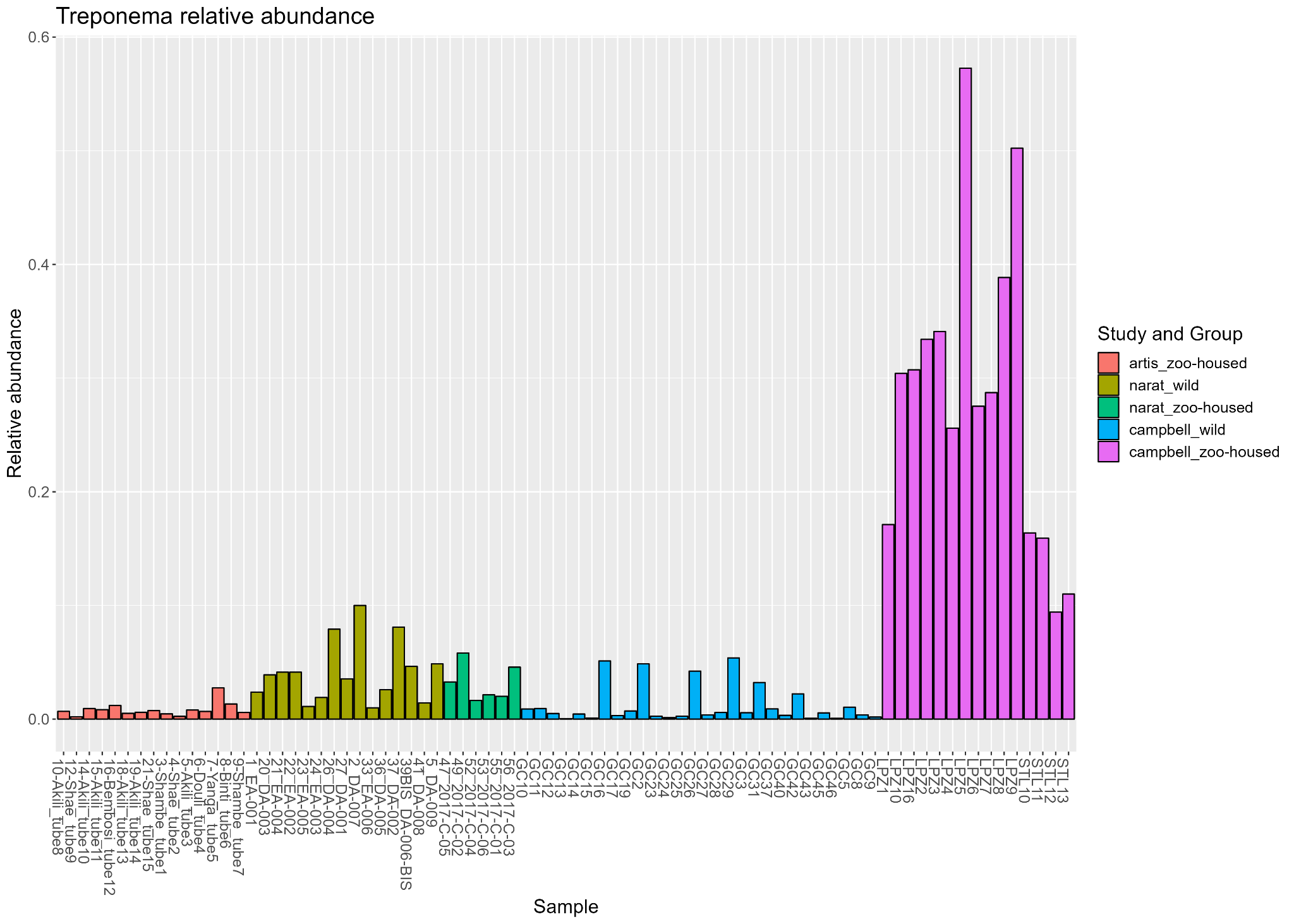

**Figure S8: Relative abundances of the genus *Treponema*.** Relative abundances (between 0 and 1) are shown per sample. Colors indicate which study a fecal sample belongs to and whether it originates from a wild or zoo-housed gorilla.

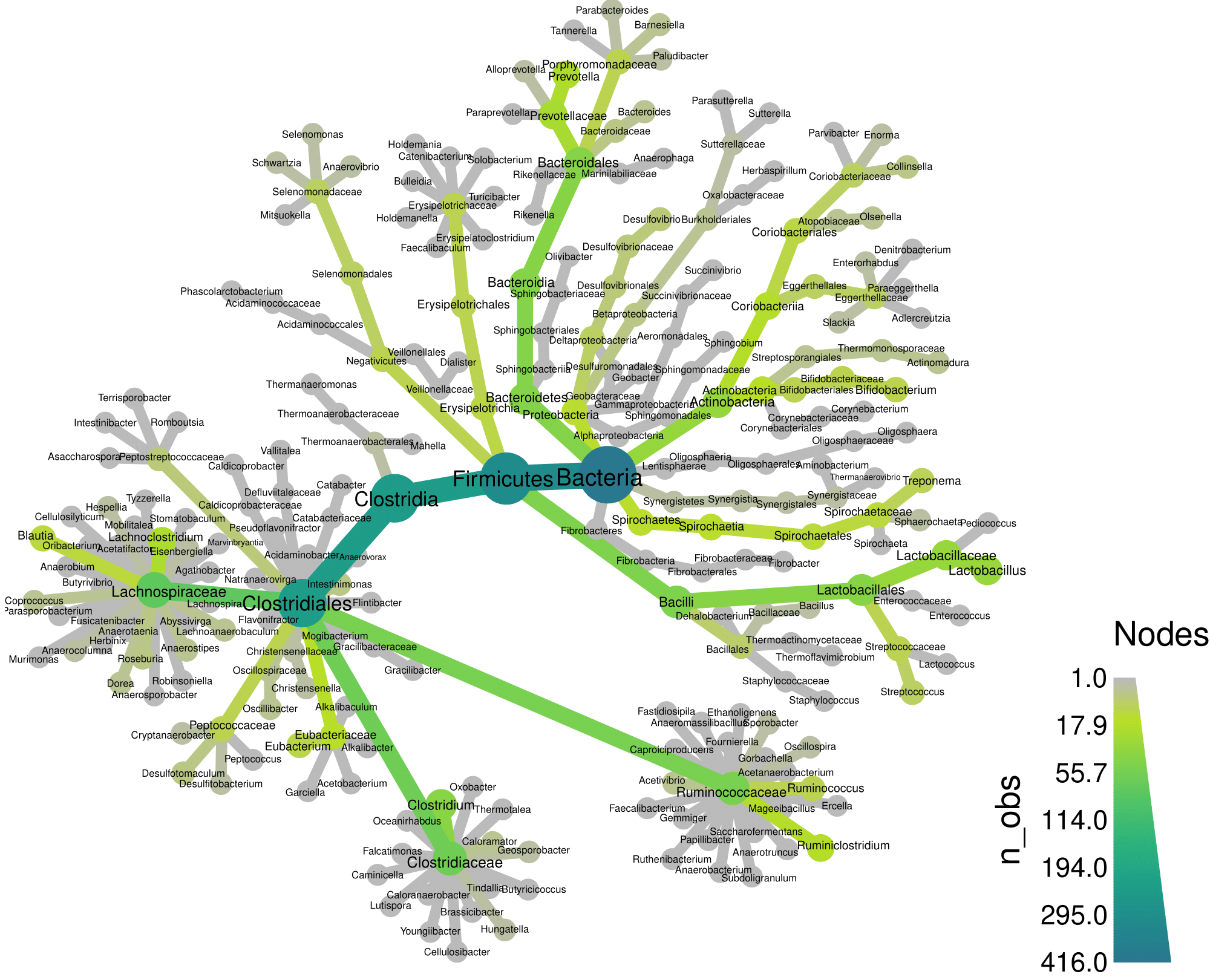

**Figure S9: Heat trees illustrating the taxonomic composition of zoo-housed ARTIS gorilla microbiota based on 16s rRNA amplicon sequencing**. Node size and color represent the number of observations at each taxonomic level.

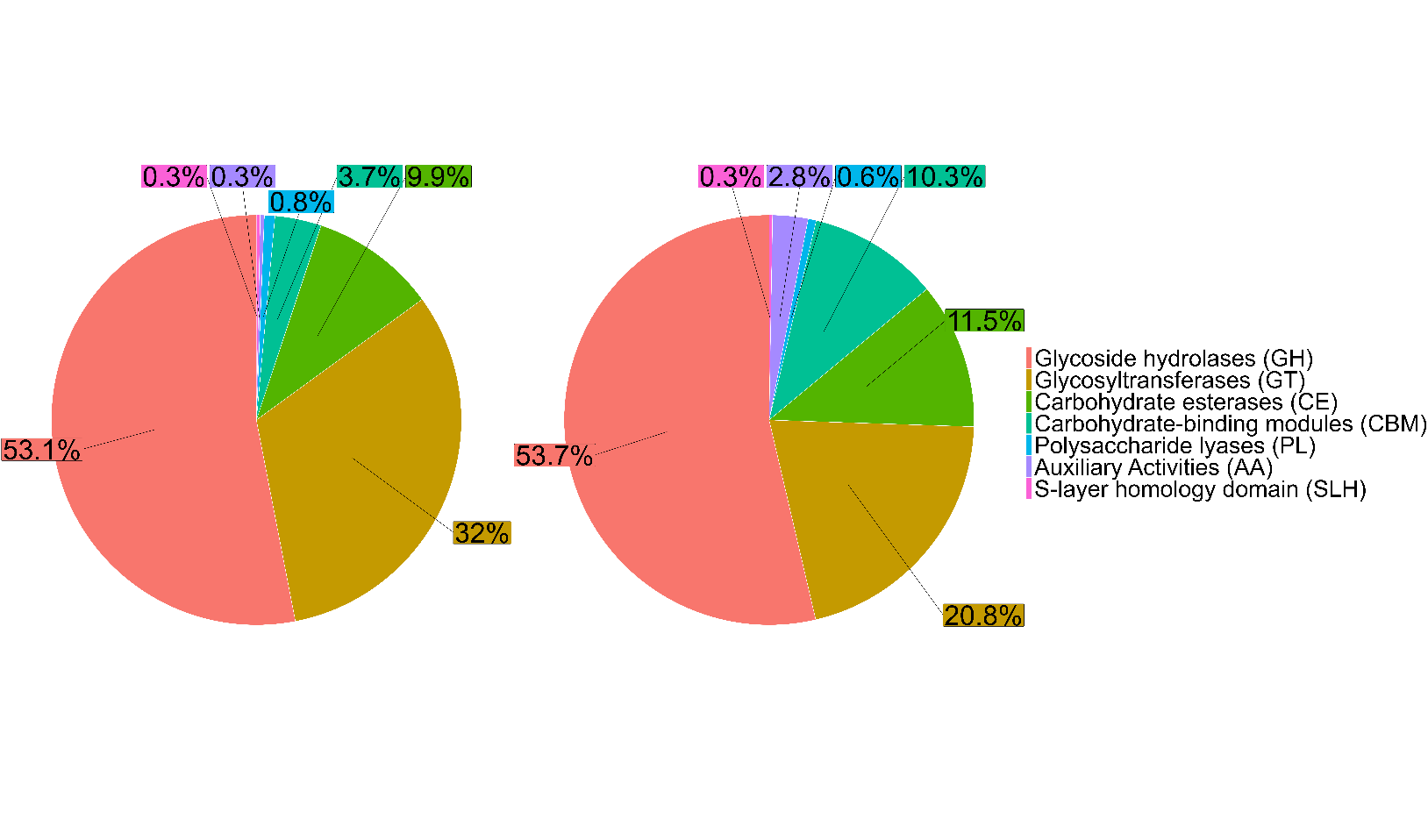

**Figure S10: Distribution of CAZyme classes in the zoo-housed gorilla microbiome (DNA and RNA).** Mean percentual contribution of CAZyme classes tot the total number of detected Carbohydrate Active Enzymes for metagenome (DNA, left) and metatranscriptome (RNA, right) of four fecal samples of a zoo-housed gorilla.

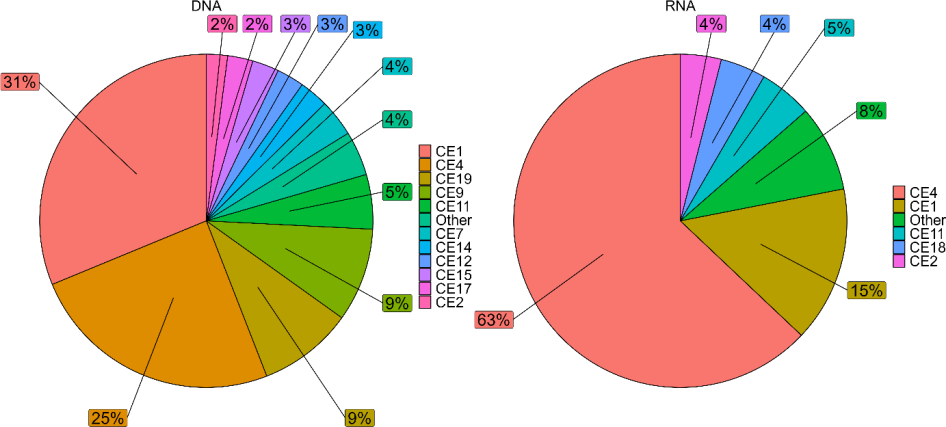

**Figure S11: CE families in the gut metagenome and metatranscriptome of a zoo-housed gorilla (DNA vs RNA).** Percentages represent the percentual contribution of each Carbohydrate esterase (CE) family to the total number of CE families found by shotgun metagenomics (DNA, left) and RNA-seq (RNA, right) on fecal samples of a zoo-housed gorilla.

**Figure S12: CBM families in the gut metagenome and metatranscriptome of a zoo-housed gorilla (DNA vs RNA).** Percentages represent the percentual contribution of each Carbohydrate Binding Module (CBM) family to the total number of CBM families found by shotgun metagenomics (DNA, left) and RNA-seq (RNA, right) on fecal samples of a zoo-housed gorilla.
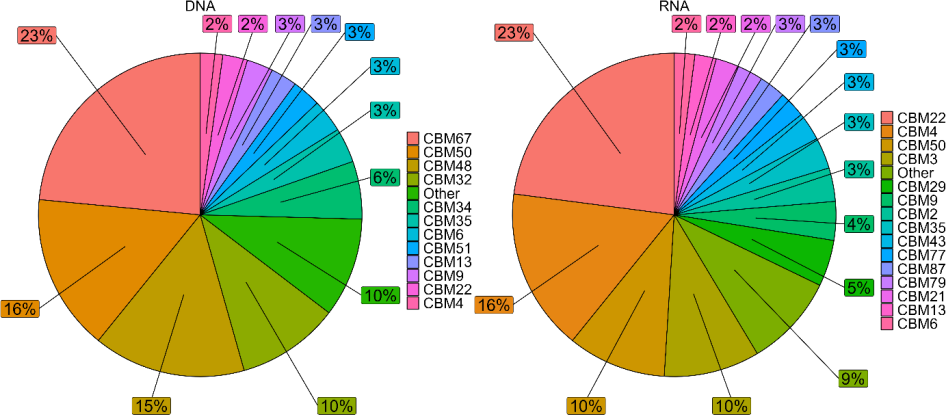

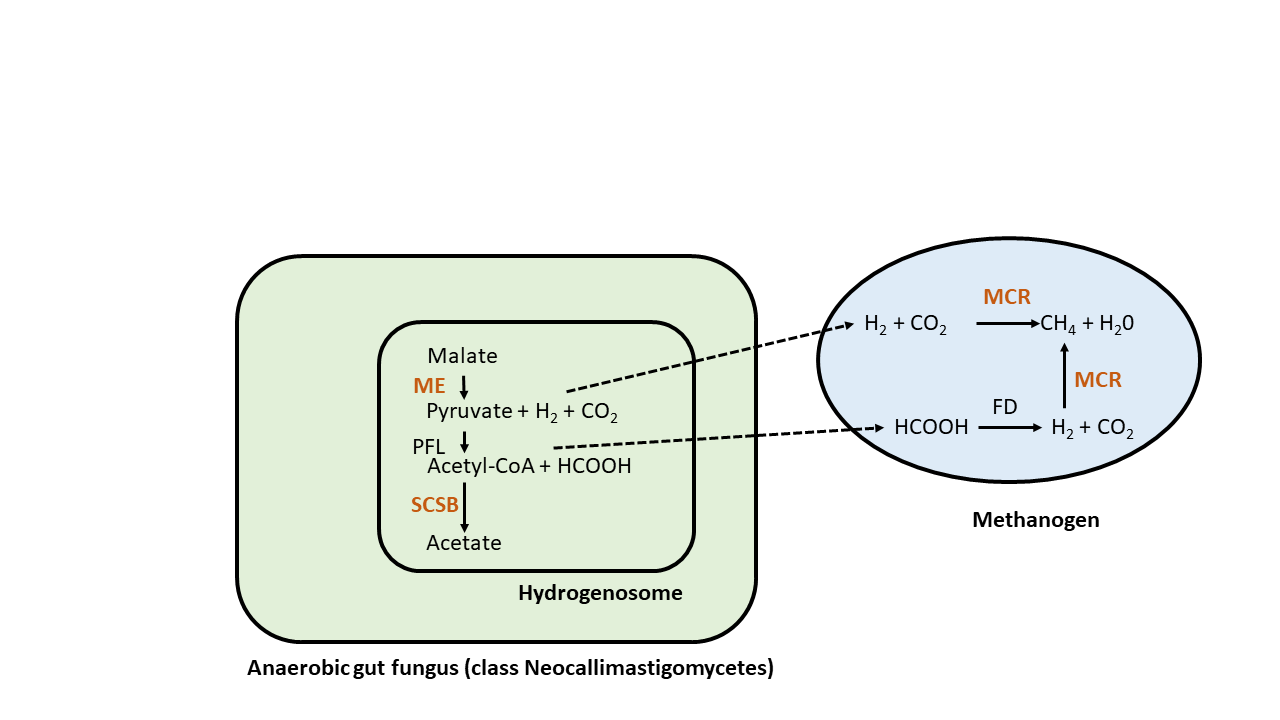

**Figure S13. Hydrogenosome-methanogenesis pathway interaction between anaerobic fungi and methanogens.** The hydrogen and formate produced by anaerobic fungi when degrading plant polysaccharides is used by methanogens to produce methane. Enzymes for which transcripts were detected are indicated in red: ME, Malic Enzyme; SCSB, Subunit B of Succinyl-CoA Synthetase; MCR, Methyl-coenzyme M reductase. Figure adapted from Li et al. (2016).

|  | **Top 10 genera in zoo- housed gorilla samples** | **Relative abundance (%)** | **Top 10 genera in wild gorilla samples** | **Relative** abundance **(%)** |
| --- | --- | --- | --- | --- |
| 1 | *Treponema* | 12.7 | *Flexilinea* | 15.0 |
| 2 | *Prevotella* | 8.7 | *Eubacterium* | 5.7 |
| 3 | *Lentimicrobium* | 7.9 | *Prevotella* | 5.6 |
| 4 | *Lactobacillus* | 6.8 | *Acinetobacter* | 4.2 |
| 5 | *Sporobacter* | 3.5 | *Parvibacter* | 4.1 |
| 6 | *Clostridium* | 3.2 | *unclassified Bacteroidales* | 3.2 |
| 7 | *Intestinimonas* | 2.6 | *Senegalimassilia* | 3.1 |
| 8 | *Ruminococcus* | 2.4 | *unclassified Eubacteriaceae* | 2.7 |
| 9 | *Parabacteroides* | 2.2 | *Clostridium* | 2.6 |
| 10 | *Oscillospira* | 2.1 | *Mogibacterium* | 2.6 |

| Phylum | Zoo-housed gorillas | Wild gorillas |
| --- | --- | --- |
| Firmicutes | 47.2% | 45.4% |
| Bacteroidetes | 30.2% | 14.9% |
| Spirochaetes | 13.4% | 2.4% |
| Actinobacteria | 4.0% | 14.2% |
| Proteobacteria | 2.7% | 6.5% |
| Chloroflexi | 0.13% | 15.1% |

**Table S1: Relative abundances of the top 10 bacterial genera in zoo-housed gorilla and wild gorilla samples.**

**Table S2: Relative abundances of the top 6 bacterial phyla in zoo-housed gorilla and wild gorilla samples based on 16s rRNA amplicon sequencing.**

|  | **Top 10 bacterial genera in zoo- housed metatranscriptome** | **Relative abundance (%)** |
| --- | --- | --- |
| 1 | *Lactobacillus* | 14.6 |
| 2 | *Limosilactobacillus* | 9.4 |
| 3 | *Muricauda* | 4.2 |
| 4 | *Clostridium* | 2.9 |
| 5 | *Prevotella* | 2.1 |
| 6 | *Treponema* | 2.1 |
| 7 | *Vibrio* | 1.6 |
| 8 | *Streptomyces* | 1.6 |
| 9 | *Ruminococcus* | 1.2 |
| 10 | *Blautia* | 0.8 |

**Table S3: Relative abundances of the top 10 bacterial genera in zoo-housed gorilla gut metatranscriptome.**

|  | **Top 10 eukaryotic genera in zoo- housed metatranscriptome** | **Relative abundance (%)** |
| --- | --- | --- |
| 1 | *Piromyces* | 9.4 |
| 2 | *Pecoramyces* | 5.3 |
| 3 | *Neocallimastix* | 4.2 |
| 4 | *Melampsora* | 0.5 |
| 5 | *Mitosporidium* | 0.3 |
| 6 | *Hirsutella* | 0.2 |
| 7 | *Aspergillus* | 0.1 |
| 8 | *Anaeromyces* | 0.1 |
| 9 | *Amorphotheca* | < 0.1 |
| 10 | *Saccharomyces* | <0.1 |

**Table S4: Relative abundances of the top 10 eukaryotic genera in zoo-housed gorilla gut metatranscriptome.**

|  | **Top 10 archaeal genera in zoo- housed metatranscriptome** | **Relative abundance (%)** |
| --- | --- | --- |
| 1 | *Methanobrevibacter* | 8.3 |
| 2 | *Candidatus Methanoplasma* | 6.8 |
| 3 | *Candidatus Methanomethylophilus* | 3.8 |
| 4 | *Methanomassiliicoccus* | 0.4 |
| 5 | *Methanobacterium* | <0.1 |
| 6 | *Methanoregula* | <0.1 |
| 7 | *Halorussus* | <0.1 |
| 8 | *Candidatus Methanoperedens* | <0.1 |
| 9 | *Cuniculiplasma* | <0.1 |
| 10 | *Candidatus Micrarchaeum* | <0.1 |

**Table S5: Relative abundances of the top 10 archaeal eukaryotic genera in zoo-housed gorilla gut metatranscriptome.**

| **Class** | **Function** |
| --- | --- |
| Glycoside hydrolases (GH) | Hydrolysis and/or transglycosylation of glycosidic bonds. |
| Glycosyltransferases (GT) | Biosynthesis of glycosidic bonds from phospho-activated sugar donors. |
| Polysaccharide lyases (PL) | Cleavage of the glycosidic bonds of uronic acid-containing polysaccharides by a β-elimination mechanism. |
| Carbohydrate esterases (CE) | Removal of ester-based modifications present in mono-, oligo- and polysaccharides, thereby facilitating the action of GHs on complex polysaccharides. |
| Carbohydrate-binding modules (CBM) | Targeted to substrate and promoting prolonged interacting, thereby potentiating the enzymatic activities of CAZymes. |
| Auxiliary Activities (AA) | Catalytic enzymes potentially involved in plant cell degradation through an ability to help the original GH, PL and CE enzymes gain access to the carbohydrates comprising the plant cell wall |

**Table S6: CAZyDB classes and their associated function.**

| **UniProt ID** | **Samples** | **Name** | **Organism** | **UniProt Annotation status** |
| --- | --- | --- | --- | --- |
| **H2BPU6** | ALL | Putative cellulase | *Neocallimastix patriciarum* (rumen fungus) | Evidence at transcript level |
| **Q8J1E3** | ALL | Cellulase Cel48A | *Piromyces sp. (strain E2)* | Evidence at transcript level |
| **A0A220DC16** | ALL | Glycoside hydrolase family 48 protein | *uncultured actinobacterium (soil)* | Predicted |
| **R9T9W1** | 15_07_2_RNA,  09_08_1_RNA,  09_08_2_RNA | Glycoside hydrolase 48 family protein | *uncultured bacterium (rumen)* | Predicted |
| **R9T9K1** | 15_07_1_RNA, 15_07_2_RNA | Glycoside hydrolase 48 family protein | *uncultured bacterium (rumen)* | Predicted |
| **R9TAI7** | 15_07_2_RNA | Glycoside hydrolase 48 family protein | *uncultured bacterium (rumen)* | Predicted |
| **R9TCD2** | ALL | Glycoside hydrolase 48 family protein | *uncultured bacterium (rumen)* | Predicted |
| **R9TDC9** | 09_08_1_RNA, 09_08_2_RNA | Glycoside hydrolase 48 family protein | *uncultured bacterium (rumen)* | Predicted |
| **B0FEW0** | 15_07_1 RNA, 15_07_2 RNA | Cellobiohydrolase | *Piromyces rhizinflatus* | Evidence at transcript level |

**Table S7: GH6 and GH48 proteins detected in zoo-housed metatranscriptome.** Protein sequences belonging to families GH6 and GH48 detected with ShortBRED amongst RNA reads originating from the ZHG microbiome.

| **UniProt ID** | **Samples** | **Name** | **Organism** | **UniProt Annotation status** |
| --- | --- | --- | --- | --- |
| **A0A1Y1WF22** | 15_07_1_RNA, 15_07_2_RNA | Homoaconitase, mitochondrial | *Anaeromyces robustus* | Unreviewed- protein inferred from homology |
| **A0A1Y1V9J2** | 15_07_2_RNA | Arg5,6 arginine biosynthetic enzyme | *Piromyces finnis* | Unreviewed- protein inferred from homology |
| **A0A1Y1WWZ0** | 15_07_1_RNA, 15_07_2_RNA,  09_08_1_RNA | Adenylate kinase | *Anaeromyces robustus* | Unreviewed- protein inferred from homology |
| **A0A1Y1XN91** | 15_07_2_RNA | Dihydroorotate dehydrogenase (quinone), mitochondrial | *Anaeromyces robustus* | Unreviewed- protein inferred from homology |
| **A0A1Y2CVY2** | 15_07_2_RNA | Aconitate hydratase, mitochondrial | *Neocallimastix californiae* | Unreviewed- protein inferred from homology |
| **A0A1Y2DRS2** | 15_07_2_RNA | NADH dehydrogenase [ubiquinone] flavoprotein 1, mitochondrial | *Neocallimastix californiae* | Unreviewed- protein inferred from homology |
| **A0A1Y1XPP5** | 15_07_1_RNA, 15_07_2_RNA | Alanine--tRNA ligase | *Anaeromyces robustus* | Unreviewed- protein inferred from homology |
| **A0A1Y2E900** | 15_07_1_RNA, 15_07_2_RNA | Dynamin-type G domain-containing protein | *Neocallimastix californiae* | Unreviewed- protein predicted |
| **A0A1Y2ETS6** | 15_07_1_RNA, 15_07_2_RNA | MSF1-domain-containing protein | *Neocallimastix californiae* | Unreviewed- protein predicted |
| **A0A1Y3NVB7** | 15_07_2_RNA | Aconitate hydratase, mitochondrial | *Piromyces sp. (strain E2)* | Unreviewed- protein inferred from homology |
| **P53587** | 15_07_1_RNA, 15_07_2_RNA | Succinate--CoA ligase [ADP-forming] subunit beta, hydrogenosomal | *Neocallimastix frontalis (Rumen fungus)* | Reviewed - Experimental evidence at transcript level |
| **Q7Z941** | ALL | Succinate--CoA ligase [ADP-forming] subunit alpha, mitochondrial | *Neocallimastix patriciarum (Rumen fungus)* | Unreviewed - Experimental evidence at transcript level |

**Table S8: Hydrogenosomal transcripts detected in ZHG gut metatranscriptome.** Hydrogenosomal or malic enzyme encoding protein sequences belonging to Neocallimastigomycetes detected with ShortBRED amongst RNA reads originating from the ZHG microbiome.
